# Inhibition of the mitochondrial permeability transition exerts sex-specific stimulatory effect on fracture repair

**DOI:** 10.1101/771691

**Authors:** Brianna H. Shares, Tzong-Jen Sheu, Kevin Schilling, Rubens Sautchuk, Ananta Paine, Gisela Beutner, Laura C. Shum, Charles O. Smith, Emma Gira, Edward Brown, Hani Awad, Roman A. Eliseev

## Abstract

Bone fracture is accompanied by mechanical stresses and inflammation – conditions that impair mitochondria via the phenomenon of permeability transition. This phenomenon occurs due to opening of the mitochondrial permeability transition pore (MPTP) promoted by cyclophilin D (CypD). MPTP opening exacerbates inflammation and cell death and, thus can disrupt fracture repair. Here we tested a hypothesis that protecting mitochondria from MPTP opening via inhibition of CypD improves fracture repair. Our data indicate that osteoblast activity, bone formation, and biomechanical properties of repaired bones were significantly increased in CypD knock-out mice when compared to controls during fracture repair. These effects were observed in male but not female mice, thus showing sexual dimorphism. Pharmacological inhibition of CypD with NIM811 in male mice also stimulated fracture repair. In addition, CypD knock-out or pharmacological inhibition produced pro-osteogenic effect in isolated bone marrow osteoprogenitors. This *in vitro* effect was associated with higher mitochondrial respiration and increased β-catenin activity regulated by mitochondria-dependent acetylation. Our findings implicate a sex-specific role of MPTP in bone fracture and suggest CypD inhibition as a modality to promote fracture repair.

## Introduction

The incidence of bone fractures is increasing worldwide as a function of the growth of the aged population over 50 years old with osteoporosis and increased bone fragility (1). Currently, 5-10 % of fractures result in fracture non-unions, a condition in which few bone-anabolic treatment modalities have proven effective (2). Fracture healing is a multistage process involving cells of various lineages including immune cells, chondro- and osteoprogenitors, endothelial cells, etc. Among them, bone marrow stromal (a.k.a. mesenchymal stem) cells (BMSCs) and periosteal osteoprogenitors are essential to the fracture healing process because these cells can differentiate into cartilage and then bone through a process of endochondral ossification or directly into bone through a process of intramembranous ossification (3). We and others have shown that differentiation of osteoprogenitors into osteoblasts (OB) involves activation of mitochondrial oxidative phosphorylation (OxPhos) (4–6). It has also been well established that in various pathological settings, including trauma, mitochondrial dysfunction is a pronounced phenotype. Mitochondrial dysfunction is even more pronounced in aging. However, the relationship between mitochondrial metabolism and fracture healing has not been studied (7).

BMSCs are characterized by their tri-lineage (osteo-, adipo-, and chondro-genic) differentiation potential, surface markers (e.g. cd31-,cd45-, cd29+, cd105+), and adherence to plastic (8). BMSCs together with periosteal cells are needed for bone maintenance during normal remodeling and for repair in response to fracture (9). Differentiation of BMSCs into OBs is an energy demanding process for which mitochondria are the most suitable sources of energy due to their highly efficient ATP production. Mitochondria also serve as major biosynthetic and signaling hubs which are very important for differentiating cells. They also supply substrates for acetylation and methylation reactions required for both post-translational protein regulation and for epigenetic reactions (10).

Mitochondrial dysfunction results in impaired OB differentiation and maturation (11). Mitochondrial dysfunction is in most cases mediated through the sudden increase in permeability of the inner mitochondrial membrane to solutes of up to 1500 Da, i.e. mitochondrial permeability transition (MPT). The MPT phenomenon is a result of prolonged opening of the mitochondrial permeability transition pore (MPTP) (12). While MPTP likely has a physiological role when opened briefly or at low conductance, i.e. < 1.2 nS with diameter of less than 3 nm (12), prolonged opening of this pore at maximal conductance leads to a loss of the inner mitochondrial membrane integrity, mitochondrial swelling, and initiation of cell death. Such a prolonged opening can be induced by increases in mitochondrial ROS, calcium, misfolded proteins, and various other excessive stresses. According to recent findings by Bernardi and others (13), the pore is formed when F_O_-ATPase complexes oligomerize in the presence of a mitochondrial peptidyl-prolyl isomerase, cyclophilin D (CypD), encoded by the nuclear *Ppif* gene. To date, CypD is the only genetically proven positive regulator of MPTP opening (14) and its genetic deletion desensitizes MPTP to opening, thereby maintaining mitochondrial membrane integrity and inhibiting the MPT phenomenon (15). The earliest known inhibitor of CypD and, thus of MPTP opening is the immunosuppressor, cyclosporine A (CsA). Its immunosuppressive function stems from its ability to bind and inhibit calcineurin (16). Non-immunosuppressive derivatives of CsA have been developed, such as NIM811, Debio025, and most recently JW47 (16–18). CypD genetic deletion or pharmacological inhibition are extensively used in studies of the heart and brain pathologies, where it has been shown to protect against injury, e.g. after ischemia-reperfusion; however, the role of CypD and MPTP in bone is poorly defined (12, 14, 19–21). We were the first to demonstrate that CypD knockout (KO) mice are protected against aging-related bone loss; but exactly how CypD KO protects bone and affects osteogenic differentiation is not known (22). Investigating CypD involvement in bone pathological situations, e.g. fracture, will generate more specific insights into its pathophysiologic function since it is known that trauma, including fracture, is associated with inflammation and oxidative stress that can induce MPTP opening (23).

In addition to increased energy demands, OB differentiation requires activation of osteogenic signaling mechanisms (24, 25). One important signaling mechanism is the canonical Wnt/β-catenin pathway. This pathway is activated during the OB commitment stage (26); and β-catenin deficiency arrests OBs in early stages of differentiation (27). We recently published a report showing that activation of mitochondria in osteoprogenitors during osteogenic differentiation promotes Wnt/β-catenin signaling through an increase in β-catenin acetylation by supplying acetyl-coenzyme A (Ac-CoA, 28). This leaves the obvious question of how CypD, which controls MPTP opening and, thus mitochondrial activity, affects osteogenic differentiation promoted by β-catenin.

In this work we test the hypothesis that protecting mitochondria from MPT through inhibition of CypD upregulates osteogenesis and promotes fracture repair. We show that when compared to control mice, CypD KO mice have accelerated fracture callus ossification, remodeling, and increased biomechanical properties of repaired bone. The effect is sex-specific and present only in male mice. Male mice treated with CypD inhibitor, NIM811, also showed enhanced biomechanical properties of repaired bones. We also examined whether inhibition of CypD/MPT exerts cell-autonomous effect on BMSCs. We observed that CypD KO BMSCs or wild-type BMSCs treated with NIM811 have an increased propensity to differentiate into OBs *in vitro*. CypD KO BMSCs also form larger ossicles in an ectopic bone formation assay. We also show an increase in mitochondrial OxPhos activity accompanied by acetylation-sensitive increase in β-catenin in CypD KO BMSCs. As more effective and specific CypD inhibitors are being developed, e.g. NIM811 and JW47 (16, 17), our findings suggest CypD/MPTP inhibition as a new avenue for bone anabolic strategies in fracture repair.

## Materials and Methods

Some methods have been previously reported (5, 28) but are briefly described again here with modifications.

### STUDY APPROVAL

Animal husbandry and experiments were performed in accordance with the Division of Laboratory Animal Medicine, University of Rochester, state and federal law, and National Institutes of Health policy. University of Rochester Institutional Animal Care and Use Committee (IACUC) specifically approved this study.

### MOUSE STRAINS

C57Bl/6J mouse strain was obtained from the Jackson Laboratory (RRID: IMSR_JAX:000664). CypD KO (Ppif^-/-^) mouse strain (originated in the Molkentin lab at Cincinnati Children’s Hospital and available from the Jackson Laboratory, RRID: IMSR_JAX:009071) was obtained from the Lab of Dr. George Porter at the University of Rochester and backcrossed 5 times onto the C57Bl/6J background prior to experiments. CypD KO and control wild-type littermates were generated by breeding mice heterozygous for CypD allele. Mice were housed at 23°C on a 12-hr light/dark cycle with free access to water and PicoLab Rodent Diet 20 (LabDiet #5053, St. Louis, MO). Mice were in group housing when applicable based on weaning. Three to four-month-old healthy, test naïve mice with an average weight of 28g were used for experiments. The assessments of all animal studies were performed in a blinded and coded manner.

### FRACTURE HEALING ASSAYS

#### Tibia fracture model

Mice were subjected to unilateral tibial fractures following anesthesia as described before (43). Briefly, right tibia was exposed and mid-diaphyseal cut was made with a scalpel to induce fracture. A 27 Ga stainless steel pin was inserted intramedullary for stabilization. Nylon sutures closed the skin. Left tibia was sham operated and served as unfractured control. CypD KO and control WT littermate mice that underwent tibial fracture were collected at post fracture day (PFD) 7, 14, 21, 28, and 35 for analyses. C57Bl/6J mice that underwent fracture were randomly divided into two groups and treated with either NIM811 (Labnetwork Inc, Cambridge, MA) or vehicle control. Freshly diluted NIM811 was injected intraperitoneally three times per week for the duration of the fracture healing (35 days) at a dose of 5 mg/kg, while control mice were injected with vehicle. Vehicle consisted of 7 parts PBS, 2 parts Kolliphor oil, and 1 part 100% EtOH. NIM811 dosage and regimen was based on previous rodent and human studies (20,29,30).

#### Biomechanical torsion testing

Following euthanasia, the fractured and unfractured tibiae were isolated and cleaned of excess soft tissue. Tibiae were stored at −80°C; and pins were removed prior to biomechanical testing. Tibiae were subjected to torsional testing. The ends of the tibias were cemented (Bosworth Company) in aluminum tube holders and tested using an EnduraTec TestBench™ system (Bose Corporation, Eden Prairie, MN). The tibiae were tested in torsion until failure at a rate of 1°/sec. The torque data were plotted against rotational deformation to determine maximum torque and torsional rigidity.

#### Serum P1NP analysis

Blood was collected, and serum prepared by centrifugation and stored in −80°C until assayed. The bone formation marker, P1NP, was assessed in serum samples using the Rat/Mouse P1NP EIA kit according to the manufacturer’s instructions. Absorbance was measured at 450 nm. Data were analyzed with a 4PL curve fit to determine concentrations in ng/mL. Eight mice per collection time point were analyzed.

#### Bone micro-computed tomography

Following euthanasia, the fractured and unfractured tibiae were isolated and cleaned of excess soft tissue. Tibiae were fixed in 10 % neutral buffered formalin (NBF) for 72 hrs and pins were removed prior to micro-computed tomography (micro-CT). Tibiae were imaged using VivaCT 40 tomograph (Scanco Medical). Scanco analysis software was utilized for volume quantification. Trabecular bone vs total volume (BV/TV), bone mineral density (BMD), and cortical thickness were determined for unfractured tibias, while callus total volume, callus BV/TV, and BMD were determined for fractured tibiae.

#### Histology

NBF-fixed tibiae were processed for histology via decalcification in EDTA for two weeks followed by paraffin embedding. Sections were cut to 5 µm in three levels of each sample, then stained with either Hematoxylin/Eosin (H&E) or Alcian Blue/Hematoxylin/Orange G (ABH/OG) or with tartrate-resistant acidic phosphatase (TRAP) counter-stained with FastGreen.

#### Histomorphometry

ABH/OG-, TRAP- or immunohistochemistry (IHC)-stained slides were scanned in an Olympus VSL20 whole slide imager at 40x magnification and evaluated with VisioPharm automated histomorphometry software. ABH/OG-stained slides were analyzed to measure the bone and cartilage area relative to total non-marrow area of the callus in the fractured tibia. Fractured cortices showing complete bony bridge vs total cortical thickness of repaired bone was measured in PFD 35 sections and expressed as percent bridging. Vessel cross-sections with morphologically defined wall and lumen containing red blood cells were counted in serial ABH/OG-stained sections of the callus at PFD 7 and 14 and expressed as a mean number of vessels per section. TRAP-stained slides were analyzed to measure the TRAP positive area relative to total area of the callus in the fractured tibia. IHC-stained slides probed for osteocalcin and β-catenin were analyzed to measure the osteocalcin or β-catenin positive area relative to total area of the callus in the fracture tibia. Three different levels were counted per mouse and averaged. There were eight mice per collection time point.

#### Collagen imaging using second harmonic generation

H&E-stained slides were imaged using multiphoton microscopy. An excitation light of 810 nm was directed and focused through a water immersion objective (20X, NA 0.95, Olympus) to the sample. Backward scattered SHG was passed through the objective, filtered via a 405 ± 15 nm bandpass filter and collected in a photomultiplier tube (Hamamatsu HC125-02). The forward propagating signal from the sample was passed through a condenser lens (NA 0.9) and filtered from excess excitation light via a 670 nm short pass dichroic mirror. SHG signal was further filtered with a bandpass filter (405 ± 15 nm) and collected in a photomultiplier tube (Hamamatsu HC125-02). Images were generated using Olympus FluoView software. Forward-to-backward (F/B) scatter ratios were calculated from the gathered images using an in-house written ImageJ macro (31).

#### Immunohistochemistry

NBF-fixed tibiae were processed as above in the Histology section. IHC was carried out using a primary osteocalcin antibody (RRID:AB_10540992) diluted 1:750 or a primary β-catenin antibody (RRID:AB_331149) diluted 1:100 followed by incubation with a biotinylated anti-rabbit IgG secondary antibody (RRID:AB_2313606) diluted 1:750 and developing using a streptavidin-HRP conjugate diluted 1:1500. The primary antibody solution was composed of 1X PBS, 3% Triton X-100 and 5% Goat Serum, while the secondary and streptavidin-HRP was made in 1X PBS. Antigen retrieval was performed using a pressure cooker and Bull’s Eye Decloaker for osteocalcin and a pressure cooker and 1mM EDTA solution for β-catenin. Detection was visualized with NovaRED Peroxidase Substrate Kit and counterstained with Hematoxylin QS.

### PRIMARY CELLS AND CELL LINES

Primary bone marrow cells were isolated from CypD KO and control WT littermates or from C57Bl/6J mice by dissection and cleaning of femurs and tibiae followed by bone marrow flushing. Cells were plated at 20 x 10^6^ per 10 cm dish in low glucose DMEM (LG-DMEM) supplied with glutamine, 10% fetal bovine serum (FBS), and 1% penicillin-streptomycin. After 24h, non-adherent cells were removed and used for osteoclastogenesis assay. Adherent cell population containing bone marrow stromal (a.k.a. mesenchymal stem) cells (BMSCs) were used for BMSC experiments.

### BMSC ASSAYS

#### BMSC culture and differentiation

BMSCs were cultured in LG-DMEM described above at 37°C, 5% CO_2_, and 5% O_2_. Cells were cultured in 5% O_2_ to mimic their *in vivo* microenvironment. We used LG-DMEM containing physiological levels of glucose (5 mM) to avoid any artifacts caused by supraphysiological levels of glucose. Media was changed every day for 3 days following BMSC isolation to rid the culture of non-adherent hematopoietic cells. Media was then changed weekly until cells reached 80% confluency, which is when the cells were plated for the appropriate experiments. Osteogenic differentiation was induced by adding β-glycerol phosphate at 2 mM and ascorbate at 50 µg/mL to LG-DMEM. Osteogenic media was changed weekly until collection. To confirm osteogenic differentiation, cells were stained for osteoblast-specific alkaline phosphate (ALP) and for mineralization using alizarin red (ARed) and also analyzed for expression of osteoblast-specific genes. To inhibit ATP citrate lyase (ACLY) and thus mitochondria-dependent β*-*catenin acetylation, we used SB204990 at a concentration of 100 µM.

#### Flow cytometry

Freshly isolated bone marrow cells were resuspended in ice-cold FACS buffer (PBS added with 2% FBS) containing a mix of surface marker-specific antibodies carrying different fluorescent tags. The following BD Biosciences antibodies were used: anti-cd45 PerCP, anti-cd31 PerCP-eFluor 710, anti-cd105 eFluor450, and anti-cd29 APC. Cells were incubated at +4°C with agitation for 30 min with the above antibodies, centrifuged in a table-top centrifuge at 7000 rpm for 1 minute, washed with 500 μL ice-cold FACS buffer, centrifuged again, and finally resuspended in 250 μL ice-cold FACS buffer for analysis. Flow cytometry was performed at the University of Rochester Core Facility using BD Biosciences LSRII flow cytometer.

#### CFU assay

Bone marrow cells were isolated as described above and plated at 20,000 cells/cm^2^. LG-DMEM media change was performed in a similar manner in that media was changed daily for 3 days following isolation and then every week thereafter. Cells were stained with Crystal Violet 14 days after initial plating to assess total fibroblastic colonies (CFU-F). Clusters of greater than 50 cells were considered a CFU.

#### RNAseq

Total RNA was isolated and processed in the University of Rochester Genomic Core. The TruSeq RNA Sample Preparation Kit V2 (Illumina) was used for library construction. The amplified libraries were hybridized to the Illumina single end flow cell and amplified using cBot (Illumina). Reads were generated and aligned to the organism-specific reference genome. An adjusted *p* < 0.05 was used to determine significance.

#### Real-time RT-PCR

Total RNA was isolated using the RNeasy kit and reverse transcribed into cDNA using the qScript cDNA synthesis kit cDNA was subjected to real time reverse transcription-PCR (RT-PCR). The primer pairs used for genes of interest are outlined in Table S1. Real time RT-PCR was performed in the RotorGene system (Qiagen) using SYBR Green. The expression of genes of interest was normalized to expression of *B2m* (β-2 microglobulin).

#### Western blot

Cells were lysed with previously described lysis buffer (5) containing protease inhibitors and subjected to 4-12% sodium dodecyl sulfate polyacrylamide gel electrophoresis (SDS-PAGE) followed by transfer to polyvinylidene difluoride (PVDF) membranes and blocking in 5% dry milk reconstituted in PBST (PBS supplemented with Tween 20 at 0.05%), as previously described (5). All antibodies were diluted in 2.5% dry milk in PBST. For β*-*catenin detection, blots were probed with mouse monoclonal β*-*catenin antibody (RRID: AB_397554) diluted 1:500 and then with horseradish peroxidase (HRP) conjugated goat anti-mouse antibody (RRID: AB_11125547) diluted 1:3000. For CypD detection, blots were probed with monoclonal CypD antibody (RRID: AB_478283) diluted 1:1000 and HRP conjugated goat anti-mouse antibody diluted 1:3000. β*-* catenin signal was developed with West Femto Substrate. To verify equal loading, blots were probed with β*-*actin antibody (RRID: AB_476697) diluted 1:2000 and HRP conjugated goat anti-mouse antibody diluted 1:5000. CypD and β*-*actin signals were developed with West Pico Substrate. Bands were measured with densitometry using ImageJ software. Signal was normalized to β*-*actin.

#### Lentiviral production and infection

Plasmid constructs TOP-GFP.mC and FOP-GFP.mC were obtained in bacterial stabs (RRID: Addgene_35491 and 35492, respectively). Each plasmid was grown overnight in LB broth containing ampicillin and isolated using Qiagen Plasmid Miniprep kit. 293T cells in 10-cm plates were transfected with 2 μg of total DNA containing 1 μg of the reporter plasmid (TOP-GFP.mC or FOP-GFP.mC), 0.1 μg of the envelope plasmid (pMDG.2, RRID: Addgene_12259) and 0.9 μg of the packaging plasmid (psPAX2 RRID: Addgene_12260) using X-tremeGENE HP DNA Transfection Reagent in OPTI-Mem media to create the lentivirus. Media was changed after 24 hrs and then collected at 48 and 72 hrs post transfection. At each collection, media obtained was spun down at 1250 rpm for 5 min in a Beckman Coulter swing bucket centrifuge at 4⁰C and passed through a 0.45 µm filter to obtain the final viral product. Optimization experiments determined that 1:2 dilution of the harvested lentivirus was safe and most efficient and thus, was used for infection. BMSCs were infected with lentivirus for 48 hours. After incubation for 48 hours, cells were subjected to SB or vehicle (PBS) treatment for 24 hours to prevent any effect SB may have on infection efficiency. Cells were collected for flow cytometry analysis and distinguished by the appearance of red (indicating infection) and green positivity (indicating β-catenin TOP-GFP reporter or non-specific control FOP-GFP reporter activity).

#### Ectopic bone formation assay

Cells were loaded onto a collagen GelFoam sponge carrier at a concentration of 2 x 10^6^/50 µL supplemented with 100 ng/mL of mouse recombinant BMP-2 and subcutaneously implanted into the backs of immunocompromised nude mice. Bone formation was assessed at 6 weeks after implantation by X-ray, histology, and histomorphometry.

### OSTEOCLASTOGENESIS

Non-adherent bone marrow cells were collected from total bone marrow isolates cultured for 24 hrs and plated at 5×10^4^ cells/well in 96-well plates in α-MEM supplemented with 15% FBS, M-CSF (10 ng/ml, R&D Systems), 1% penicillin-streptomycin (Gibco), 1% non-essential amino acids (Gibco). Cells were cultured for 3 days. Thereafter, to generate osteoclasts (OCL), media was changed; M-CSF (10 ng/ml) and RANKL (10 ng/ml, R&D Systems) were added; and then media was replaced every 2 days with fresh media supplemented with fresh M-CSF and RANKL. After 5 days of RANKL exposure, cells were fixed and stained for tartrate-resistant acid phosphatase (TRAP, Sigma-Aldrich Co, 387A). OCLs were identified as multinucleated (more than 3 nuclei) TRAP-positive cells.

### MITOCHONDRIAL ASSAYS

#### Calcium Retention Capacity

Calcium Retention Capacity (CRC) assay was performed to measure MPTP opening in permeabilized cells exposed to calcium pulses as described before (13), with several modifications. In particular, BMSCs were permeabilized with 0.01% digitonin for 5 min on ice in a KCl-based buffer (140 mM KCl, 2 mM KH_2_PO_4_, 6 mM HEPES, pH 7.4) containing EGTA at 1 mM. Digitonin and EGTA were then washed away. Cells remained permeabilized even after digitonin washout as evident from positive staining of cells with Trypan Blue (not shown). One hundred thousand permeabilized cells were placed in 0.1 mL of KCl-based buffer containing 5 mM Succinate, 1 μM cytochrome c, and 1 μM Oregon Green BAPTA2, a fluorescent probe that measures extramitochondrial Ca^2+^. Of note, we used Oregon Green instead of Calcium Green 5N conventionally used in CRC assay, because of more appropriate Kd value for Ca^2+^ (0.5 μM vs 14 μM). Permeabilized cells contain significantly smaller number of mitochondria when compared to highly concentrated suspensions of isolated mitochondria usually used in the CRC assay. Therefore, we had to use smaller pulses of Ca^2+^ (0.5 μM instead of 20 μM) and higher affinity indicator, Oregon Green. Permeabilized cells were exposed to Ca^2+^ pulses, and Oregon Green fluorescence was monitored using a BioTek plate reader. Because of our plate reader specifications, we could not monitor the spikes of Ca^2+^ followed by a decrease due to uptake by mitochondria observed in previous publications (13), but could only detect a steady-state signal achieved after the pulse. MPTP opening was detected as an irreversible increase in Oregon Green fluorescence after the pulse. CRC values were expressed in nmoles of accumulated Ca^2+^ per 10^6^ cells.

#### Mitochondrial mass assay

Cells were stained with nonyl acridine orange (NAO) at 100 nM, a fluorescent probe that labels cardiolipin present primarily in mitochondrial membranes (32), for 15 min at 37°C. Stained cells were then lifted from plates with cell scraper, washed, and resuspended in PBS. NAO signal was detected in BD Biosciences LSRII flow cytometer.

#### In-gel activity of the mitochondrial respiratory complex I

Cells were lysed with lauryl maltoside at 4 µg/µg of protein and subjected to 4-10% gradient clear native polyacrylamide gel electrophoresis (CN PAGE). Complex I activity was determined as described before (33) by incubating gel in a buffer containing Tris-HCl at 5 mM (pH 7.4), NADH at 0.1 mg/mL, and nitroblue tetrazolium at 2.5 mg/mL and rocking overnight at room temperature. Complex I monomer bands at 669 kDa were measured with densitometry using Image J software. Complex I signal was normalized to VDAC signal detected by western blot done in parallel on the same samples using mouse monoclonal VDAC antibody (Abcam) diluted 1:2000 and HRP conjugated goat anti-mouse secondary antibody diluted 1:3000.

#### Seahorse metabolic profiling

Oxygen consumption rate (OCR) was measured using Seahorse XF96 metabolic profiler (Seahorse Bioscience). Cells were plated on Seahorse 96-well plates 48 hrs before experiment at a density of 30,000 cells/well. Immediately before the experiment, media was replaced with unbuffered (pH 7.4) DMEM-XF media containing glucose at 5 mM, glutamine at 1 mM, and pyruvate at 1 mM. A baseline measurement of OCR was taken and then an inhibitory analysis was performed using injections of oligomycin (Olig) at 1 µM, FCCP at 0.5 µM, and antimycin A (AntA) at 1 µM. Various OxPhos parameters were determined based on this inhibitory analysis per manufacturer’s instructions and as described in our previous work (4).

### STATISTICAL ANALYSIS

Three to eight independent experiments were done to derive each panel of the paper, depending on the results of power analysis. Power analysis was carried out using the formula: 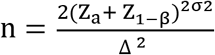, where n = sample size, α = type I error, β = type II error, Δ = effect size, σ = standard deviation, and Z is a constant (34). An example of power analysis carried out for biomechanical testing is shown below in Table S2. Data were analyzed using Prism 8 (GraphPad Software). Mean values and standard errors were calculated and the statistical significance (*p* <0.05) was established using either Student’s *t*-test when two variables were compared or one-way analysis of variance (ANOVA) when more than two variables were compared based on normal spread of our data.

## Results

### Genetic deletion of CypD leads to more efficient bone fracture repair in male mice

As fracture is associated with tissue mechanical damage and inflammation known to induce mitochondrial dysfunction via CypD-mediated opening of the MPTP, we hypothesized that protecting mitochondria via CypD deletion would have a positive effect on fracture healing. To test this, CypD KO and WT control littermates were subjected to unilateral tibial fractures; and fractured and contralateral unfractured bones were collected at various time-points to access bone fracture healing as outlined in the diagram in Figure 1A. Representative X-ray images of fractured bones at post-fracture day (PFD) 0 and 35 shown in Figure 1B demonstrate both genotypes were capable of radiographically complete healing in our time frame (stabilized tibial fracture healing in mice takes approximately 35 days (3) after which the fracture is considered biomechanically healed). We next assessed the quality of repaired bone via the biomechanical torsion test of both fractured and unfractured bones at PFD 35 in both male and female littermates. Torsional biomechanical properties in unfractured CypD KO and WT bones showed no significant differences (Supplementary Fig. S1A&B). This is in agreement with our previous observations showing that young CypD KO and WT mice have a similar skeletal phenotype at baseline (4) and that protective effects of CypD KO ensue mostly under a pathological stress, e.g. aging or trauma. Consistent with this notion, CypD KO mice showed significantly increased torsional biomechanical properties (torsional rigidity and maximum torque) of repaired tibiae when compared to WT controls (Fig. 1C, top panels); however, this difference was only observed in males, with female fractures showing no change in biomechanical properties (Fig. 1C, bottom panels). These data indicate that male CypD KO mice achieve stronger and more rigid bone repairs when compared to WT control littermates. To our knowledge this is the first report of a sexually dimorphic role of CypD/MPTP in fracture repair. Given this observation, our further efforts included only male mice in order to better characterize the role of CypD in fracture repair.

**Figure 1.**
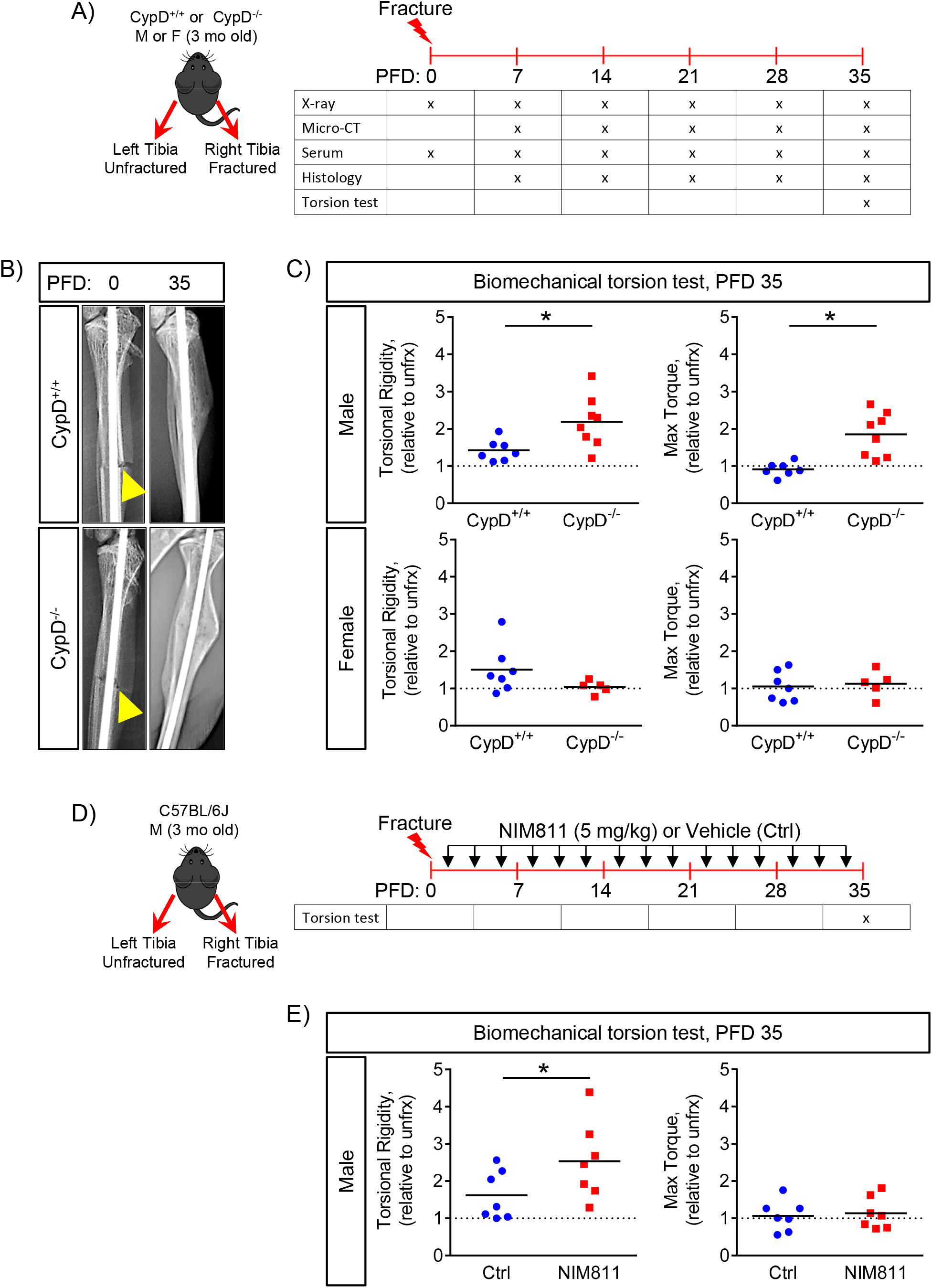
Genetic deletion or pharmacological inhibition of CypD results in stronger repaired bone in male mice at post fracture day 35. A) Tibial fractures were performed on 3-month-old male and female mice; and fractured and contralateral unfractured bones were collected for various analyses; B) Representative X-rays at post fracture day (PFD) 0 and 35. Arrowheads indicate the fracture site; C) Biomechanical properties of repaired bone were measured at PFD 35 by torsion test and reported as torsional rigidity indicating bone toughness and maximum torque indicating bone strength. Data are expressed as relative to the contralateral unfractured limb to account for differences in bone phenotype between animals. CypD^-/-^ male mice show increased torsional properties of repaired bone; whereas, CypD^-/-^ female mice show no difference when compared to control CypD^+/+^ littermates; D) Tibial fracture was induced similarly to the CypD KO mice and underwent biomechanical torsion test at PFD 35. NIM811 at 5 mg/kg or vehicle was injected intraperitoneally 3 times per week throughout the healing process; E) NIM811-treated mice show an increase in torsional rigidity and no difference in maximum torque when compared to vehicle-treated controls. Plots show actual data points and calculated means (n= 5-8). *, *p* < 0.05 vs CypD^+/+^ controls as determined by an unpaired *t*-test. See also Supplementary Figure S1 – S3.

### Pharmacologic inhibition of CypD leads to more efficient bone fracture repair in male mice

A more clinically relevant approach for CypD inhibition is using a pharmacological inhibitor. We, therefore used the non-immunosuppressive derivative of CsA, NIM811 (20,29,30). Mice that were treated with NIM811 three times per week throughout the duration of fracture repair (Fig. 1D) showed an increase in torsional rigidity when compared to vehicle-treated mice (Fig. 1E, left panel), but no difference in maximum torque (Fig.1E, right panel). Again, the unfractured tibia showed no baseline difference in biomechanical parameters (Supplementary Fig. S2). This indicates that a clinically relevant pharmacological inhibition of CypD improves bone fracture repair. The effect of the inhibitor was not as strong as CypD deletion but was nevertheless promising. Further dose and regimen optimization as well as developing bone-targeted variants may enhance the inhibitor efficacy.

**Figure 2.**
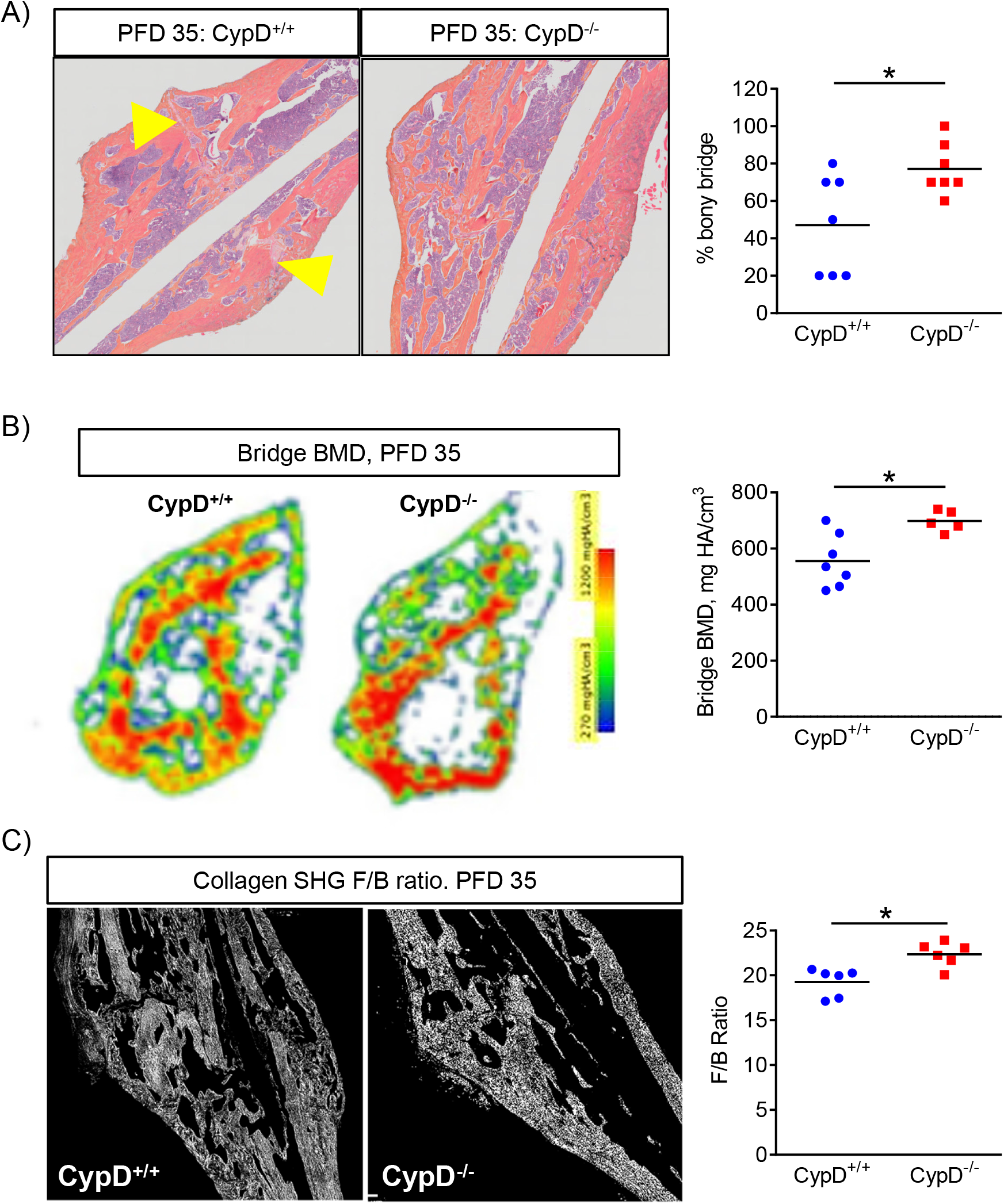
Genetic deletion of CypD results in better bone bridging at post fracture day 35. A) Representative histological sagittal sections through the callus and histomorphometric assessment of bony bridging in histological sections through the callus at PFD 35. CypD^-/-^ mice display increased bridging of broken bone; B) Representative heat map of BMD of a bridge region in a transverse section. At PFD 35, CypD^-/-^ mice show an increase in BMD at the bridge; C) Representative second harmonic generation (SHG) images of collagen in sagittal sections through the callus where forward to backward (F/B) scattering was measured using multiphoton microscopy at PFD 35 as described in Methods. CypD^-/-^ mice have higher F/B ratios indicating better aligned and more mature collagen fibers in the callus. Plots show actual data points and calculated means (n= 6-8). *, *p* < 0.05 vs CypD^+/+^ controls as determined by an unpaired *t*-test.

### CypD KO mice show increased mineralization and greater collagen matrix maturation following fracture repair

One potential explanation for increased biomechanical properties in bone repair is improved bridging of the cortical ends of fractured bone. To determine whether the increased strength and rigidity of the repaired bone in CypD KO male mice is due to differences in bridging of the cortical ends of the fractured bone, we examined histology sections of the callus at PFD 35 (Fig. 2A). We observed significant differences in bridging of the fractured bone with higher percentage of complete bridging in CypD KO mice (Fig. 2A). Since the torsional strength of the repaired bone is dependent on the mineralization of the callus bridging to reestablish continuity of the cortical ends of the fractured bone, we performed transverse section by section analysis of μCT scans and heat-mapping of BMD in sections flanking the bridge. This enabled quantification of mineralization exactly at the bony callus bridge (Fig.2B). As evident from this analysis, BMD of the bony bridge was significantly higher in CypD KO vs WT control mice (Fig. 2B, right panel). The strength and rigidity of the repaired bone is also influenced by the organic component of bone matrix, e.g. collagen I. We therefore utilized an optical method based on measuring the ratio of forward-to-backward (F/B) scattering of second harmonic generation (SHG) signal from collagen (35). Changes in F/B reflect differences in the diameter of the fibrils bundled into collagen fibers, as well as their spacing within the fiber, and their packing disorder (36). More mature collagen fibers have increased forward propagating SHG signal and ultimately have a higher F/B ratio compared to less mature collagen (37). F/B analyses of H&E-stained sections of the calluses at PFD 35 showed higher collagen F/B ratios in repaired bone of CypD KO mice when compared to WT controls (Fig. 2C). Thus, CypD KO mice have increased overall mature bone formation at the fracture site.

In addition, we performed whole callus μCT analysis at various time-points, which demonstrated with few exceptions (e.g. higher callus BMD in CypD KO vs WT mice at PFD 28) no significant differences between the CypD KO mice and WT controls in the bony callus volume fraction (BV/TV), cortical thickness, and the whole callus BMD (Supplementary Fig. S3). These data indicate that there was increased formation and higher mineralization of the bony bridge along with presence of more mature unidirectional collagen fibers in fracture calluses of CypD KO mice which together most likely accounted for improved biomechanical properties of the repaired bone.

**Figure 3.**
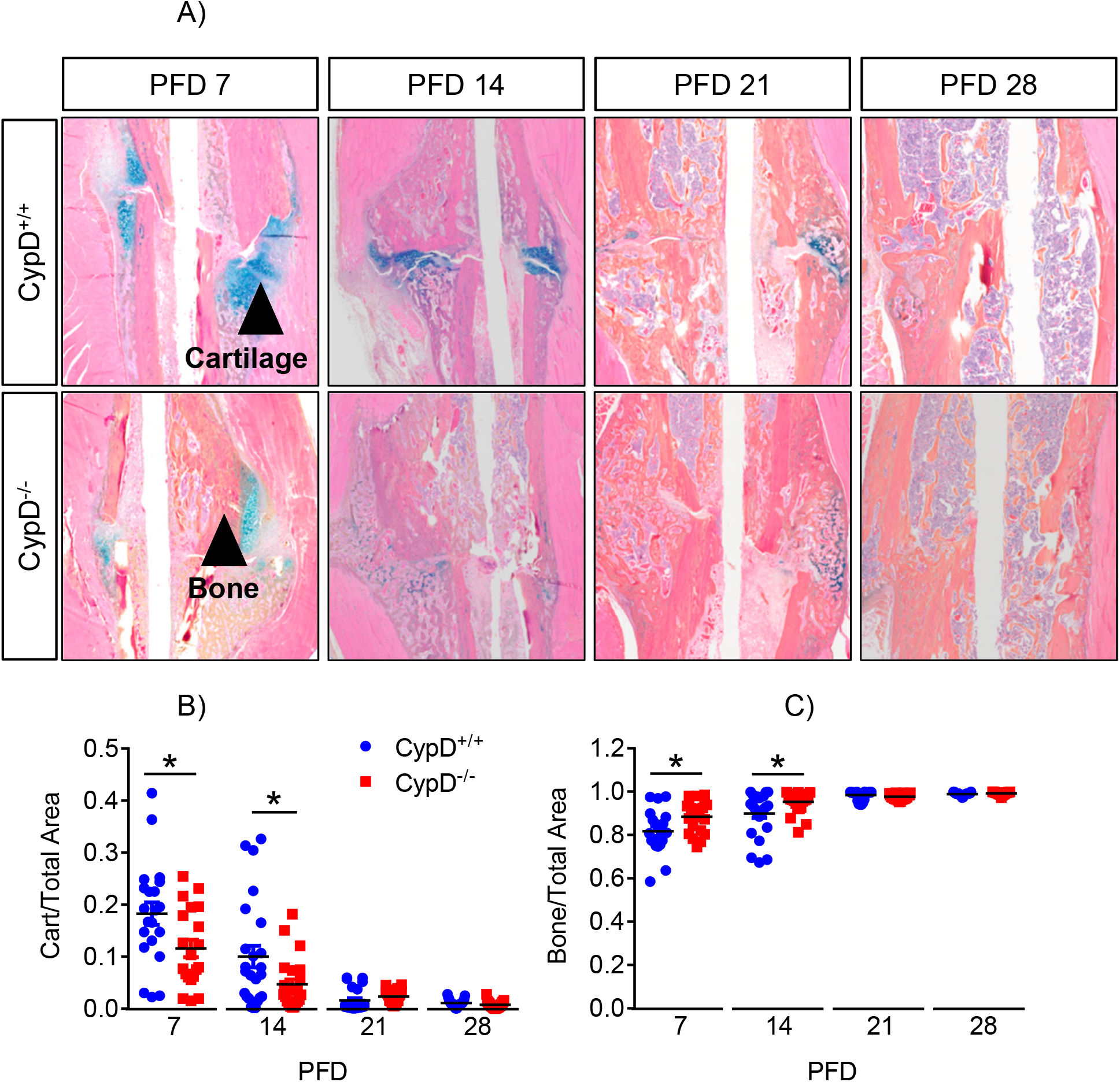
Accelerated callus ossification in CypD knock-out mice. Histological staining using ABH/OG stain followed by histomorphometry of fractured bones was performed at various time points (A). CypD^-/-^ mice show accelerated callus ossification and a truncated cartilaginous phase at PFD 7 and 14 (B & C). Plots show actual data points and calculated means (n=8, 3 replicates each). *, *p* < 0.05, determined by unpaired *t*-test of each time point. Unpaired *t*-test was used because samples at each time point were collected and analyzed independently of other time points.

### Genetic deletion of CypD results in accelerated callus ossification and remodeling during fracture healing

As our data suggest that CypD KO improves the quality of healed bone, we asked whether this was due to changes in callus composition and bone remodeling during fracture healing. Histologic analyses and automated histomorphometry of ABH/OG stained sections of the calluses of CypD KO mice at PFD 7 and 14 detected less cartilage and more bone, suggesting a truncated cartilaginous callus phase and accelerated bone formation (Fig. 3A-C). Immunostaining for osteoblastic marker, osteocalcin, in CypD KO mice revealed an increase in osteocalcin within the callus at PFD 7 and again at PFD 28 when compared to WT controls (Fig. 4A), supporting the notion of accelerated bone formation in CypD KO fracture calluses seen in Figure 3. To further confirm the effect on bone formation, we measured serum P1NP, a procollagen peptide synthesized by OBs during bone formation process (26). When compared to WT control littermates, CypD KO mice show an increase in serum P1NP levels at PFD 7 which did not reach statistical significance (*p* = 0.2873) and statistically significant increases at PFD 28 and 35 (Fig. 4B). Together, increased osteogenic markers at PFD 7 likely indicate accelerated new bone formation while at PFD 28 and 35 accelerated bony callus remodeling during fracture repair in CypD KO mice.

**Figure 4.**
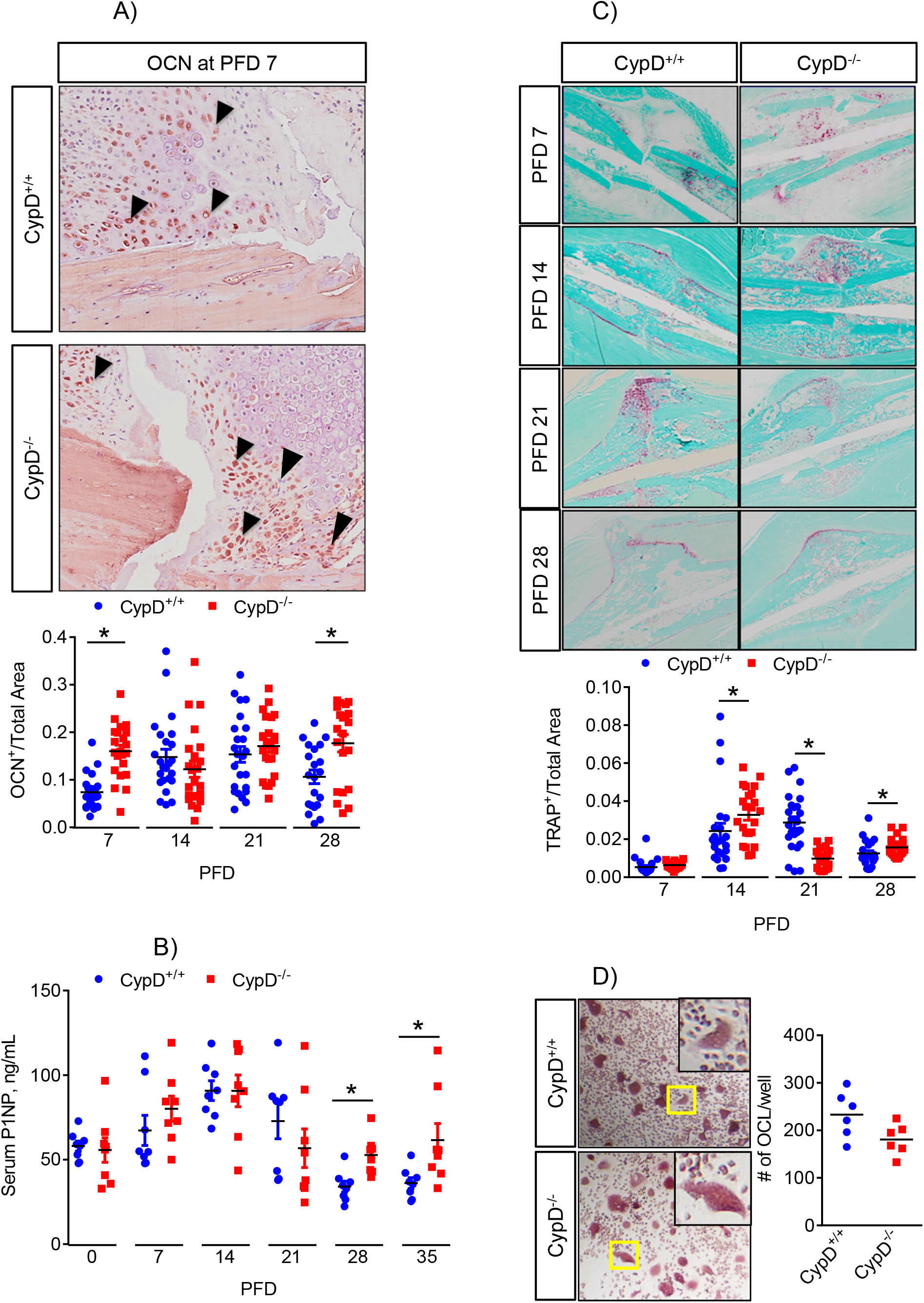
Genetic deletion of CypD results in increased bone formation and remodeling during fracture healing process. Sections of calluses were probed with antibodies against osteocalcin (OCN) at various time-points (A). CypD^-/-^ mice show increased OCN immunostaining at PFD 7 and 28. B) Serum was collected at various time-points and analyzed for P1NP. CypD^-/-^ mice show increased serum P1NP at PFD 28 and 35 when compared to CypD^+/+^ control littermates, indicating an increase in bone formation. Tissue sections were stained with TRAP-specific stain to detect osteoclast activity (C). CypD^-/-^ mice show increased TRAP staining and, thus bone remodeling at PFD 14 and again at PFD 28; D) Osteoclast (OCL) precursors were isolated from bone marrow and OCL differentiation assay was performed as described in Methods. Number of OCL per well was not significantly different between CypD^-/-^ and CypD^+/+^ control littermates. Boxed regions in (D) are magnified in the inserts to show details of multinucleated TRAP^+^ cells. Plots show actual data points and calculated means. *, *p* < 0.05, determined by unpaired *t*-test of each time point. Unpaired *t*-test was used because samples at each time point were collected and analyzed independently of other time points. See also Supplementary Figures S4 & S5.

Accelerated remodeling of bony callus would suggest an increase in osteoclasts at the callus. Osteoclast-specific TRAP staining revealed that CypD KO mice remodeled callus more actively than control littermates at both PFD 14 and 28 (Fig. 4C). TRAP staining also peaked earlier in CypD KO when compared to WT samples: at PFD 14 vs PFD 21, respectively, consistent with the hypothesis of increased remodeling in the newly formed bone. Increased levels of osteocalcin and P1NP (Fig. 4A & B) observed in CypD KO mice at later stages of fracture repair are consistent with more remodeling at this stage and confirmed by higher TRAP staining in CypD KO mice. To find out whether CypD KO had a direct effect on osteoclasts (OCL), we performed OCL differentiation assay, which showed that OCL differentiation was not significantly different between CypD KO and control (Fig. 4D). OCLs are regulated by BMSC- and OB-derived RANKL and Opg and, in particular, by RANKL/Opg ratio. We, therefore examined expression of their corresponding genes, *Tnfsf11* and *Tnfrsf11b* using our RNAseq data. Our data showed that although the expression of both genes was downregulated in CypD KO BMSCs when compared to controls, their expression ratio was unchanged (Supplementary Fig. S5A). These data indicate that OCL may play a role in the accelerated healing phenotype seen in CypD KO, but this role is most likely secondary to the OB role in this process.

As fracture callus ossification and, therefore remodeling may be influenced by changes in callus vascularization, we counted the number of blood vessels in calluses at PFD 7 and 14 using histomorphometry. There were no significant differences in the number of vessels between CypD KO and WT control littermates (Supplementary Fig. S4). It is, therefore unlikely that the observed changes in the callus ossification was due to changes in vascularization. Together, these histological and serological analyses indicate that CypD KO mice have an accelerated callus ossification and remodeling with truncated cartilaginous phase during fracture healing and these effects are not accompanied by changes in callus vascularization.

### Genetic deletion or pharmacological inhibition of CypD enhances osteogenic function of BMSCs

BMSCs together with periosteal progenitors are precursors of OBs that are responsible for new bone formation during fracture repair. Since we observed accelerated bone formation during fracture repair in CypD KO mice, we investigated whether there is a cell-autonomous pro-osteogenic effect of CypD KO in BMSCs. We first investigated whether CypD KO affected BMSC population using two general methods to assess bone marrow isolates: detection of surface markers by flow cytometry in freshly isolated bone marrow and colony forming unit (CFU) assay. We found that CypD KO did not cause any significant shifts in either the size of cd45-cd31-cd105+ cd29+ population or CFUs (Supplementary Fig. S5B & C). Next, we performed *in vitro* osteoinduction as well as *in vivo* ectopic bone formation assays. CypD KO BMSCs showed a more pronounced increase both in alkaline phosphatase (ALP, Fig. 5A) and alizarin red (ARed, Fig. 5B) staining when compared to WT control cells after 14 days of osteoinduction. Real-time RT-PCR analysis of expression of osteogenic marker genes during osteogenic differentiation showed that CypD KO BMSCs have similar expression of bone sialoprotein (*Ibsp*) but significantly higher expression of osteocalcin (*Bglap*) when compared to the WT cells (Fig. 5C) at both Day 0 and Day 14. This variation in the effect on expression of these two osteogenic markers may be explained by the difference in gene regulation. For example, it is known that *Bglap* is a direct transcriptional target of TCF/β-catenin while *Ibsp* is not (38). Altogether, these data indicate that BMSC from CypD KO mice have an increased osteogenic potential.

**Figure 5.**
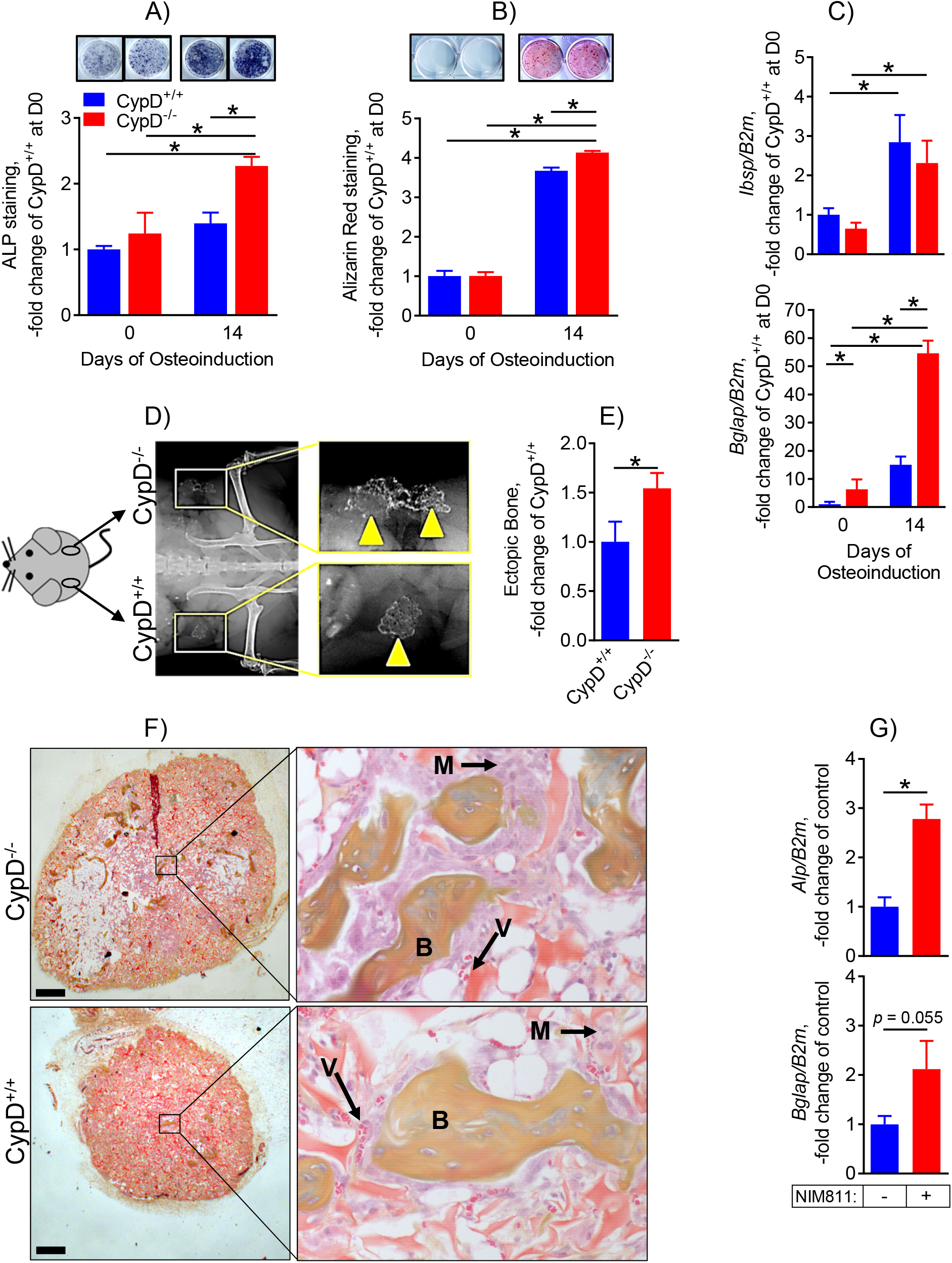
CypD knock-out BMSCs show increased osteogenic potential. BMSCs from CypD^-/-^ and control CypD^+/+^ littermates were osteoinduced for 0 or 14 days and analyzed by staining for alkaline phosphatase (ALP, A) and alizarin red (ARed, B) and by real-time RT-PCR (C). CypD^-/-^ BMSCS show more intense ALP and ARed staining (A&B) and higher expression of osteogenic gene marker, *Bglap* (C). Data are presented as fold changes over undifferentiated control BMSCs. BMSCs were also implanted into backs of immunocompromised nude mice for an ectopic bone formation assay (D – F). Ossicle formation was analyzed after 6 weeks; D) X-ray showing ectopic bone formation; E) Ectopic bone was measured with histomorphometry; F) Histology sections of ectopic bones with boxed areas magnified showing bone (B), marrow (M), and blood vessels (V). CypD^-/-^ BMSCs show a significant increase in ectopic bone formation as analyzed with histomorphometry; G) BMSCs isolated from C57BL/6J mice were induced in osteogenic media for 14 days in the presence of NIM811 at 0.5 mM or vehicle control; and osteogenic marker gene expression was measured with real-time RT-PCR. NIM811-treated cells had significantly higher expression of *Alp.* Data are means ± SEM (n=3-6). *, *p* < 0.05, determined by ANOVA followed by post-hoc analysis and unpaired *t*-test where applicable based on specifications in Methods. See also Supplementary Figure S5.

In order to demonstrate the increased osteogenic potential of CypD KO BMSC *in vivo*, we performed BMP2-mediated ectopic bone (ossicle) formation assay. The assay is based on subcutaneous implantation of cells with carriers (Gelfoam^®^ collagen sponges soaked in BMP2) in the back of immunocompromised nude mice as previously described (39). Grafted BMSC are known to create bone and support both development of vascularized bone marrow of recipient origin and, according to some reports, osteoinduction in host fibroblasts (39). At 6 weeks following implantation of both WT (left flank) and CypD KO BMSCs (right flank), we were able to confirm ossicle formation in grafts from both genotypes (Fig. 5D). CypD KO BMSCs grafts led to significantly larger ossicles as assessed with histomorphometry (Fig. 5E). Importantly, both CypD KO and WT BMSCs grafts led to formation of ectopic bone, marrow, and vascularized stroma (Fig. 5F). Together with the *in vitro* results, these data confirm that CypD KO BMSCs have an increased osteogenic potential. We also tested whether pharmacological inhibition of CypD with NIM811 would exert similar pro-osteogenic effects. Figure 5G shows that osteoinduction of BMSCs in the presence of NIM811 for 14 days increased osteogenic markers *Alpl* and *Bglap* when compared to controls (Fig. 5G). As all these effects are observed in isolated cells that are deprived of systemic influences present *in vivo*, the most likely explanation for the pro-osteogenic effect of CypD deletion or pharmacological inhibition on BMSCs is a cell-autonomous mechanism.

### Genetic deletion of CypD leads to inhibition of MPTP opening and increased mitochondrial activity in BMSCs

CypD KO desensitizes mitochondria to MPTP opening (14, 40), protects mitochondria from various stresses, and improves OxPhos likely via better assembly of mitochondrial respiratory complexes (41). Previous work from our lab and others indicates that increased mitochondrial OxPhos stimulates osteogenic signaling in primary BMSCs and osteogenic cell lines (4,5,42). We, therefore investigated whether the observed pro-osteogenic effect of CypD KO in BMSCs is associated with improved OxPhos. Our data show that when compared to WT controls, CypD KO BMSCs show higher mitochondrial Calcium Retention Capacity (CRC), indicative of decreased MPTP activity. Mitochondria in CypD KO BMSCs retain higher amounts of calcium (7 vs 3 nmoles Ca^2+^/10^6^ cells) before undergoing permeability transition (Fig. 6A). To confirm that this higher CRC was not due to higher number of mitochondria in CypD KO cells, we measured mitochondrial mass in WT and CypD KO BMSCs with nonyl acridine orange (NAO) and flow cytometry (Supplementary Fig. S6A). The assay showed no significant difference in NAO signal and, thus in mitochondrial mass between WT and CypD KO cells. As decreased MPTP activity is linked to increased mitochondrial efficiency likely via better assembly and function of the mitochondrial respiratory Complex I (CxI, 41), we measured CxI activity using native gel electrophoresis and In-gel colorimetric assay. CypD KO BMSCs displayed higher CxI activity than the WT cells (Figures 6B & C). Interestingly, CxI activity increased in both WT and CypD KO cells during osteoinduction, so that the difference between them was no longer significant (Supplementary Fig. S6B). This suggests a mechanism whereby CypD KO mitochondria are maintained at near maximal capacity at the undifferentiated stage, allowing for the increased osteogenic differentiation.

**Figure 6.**
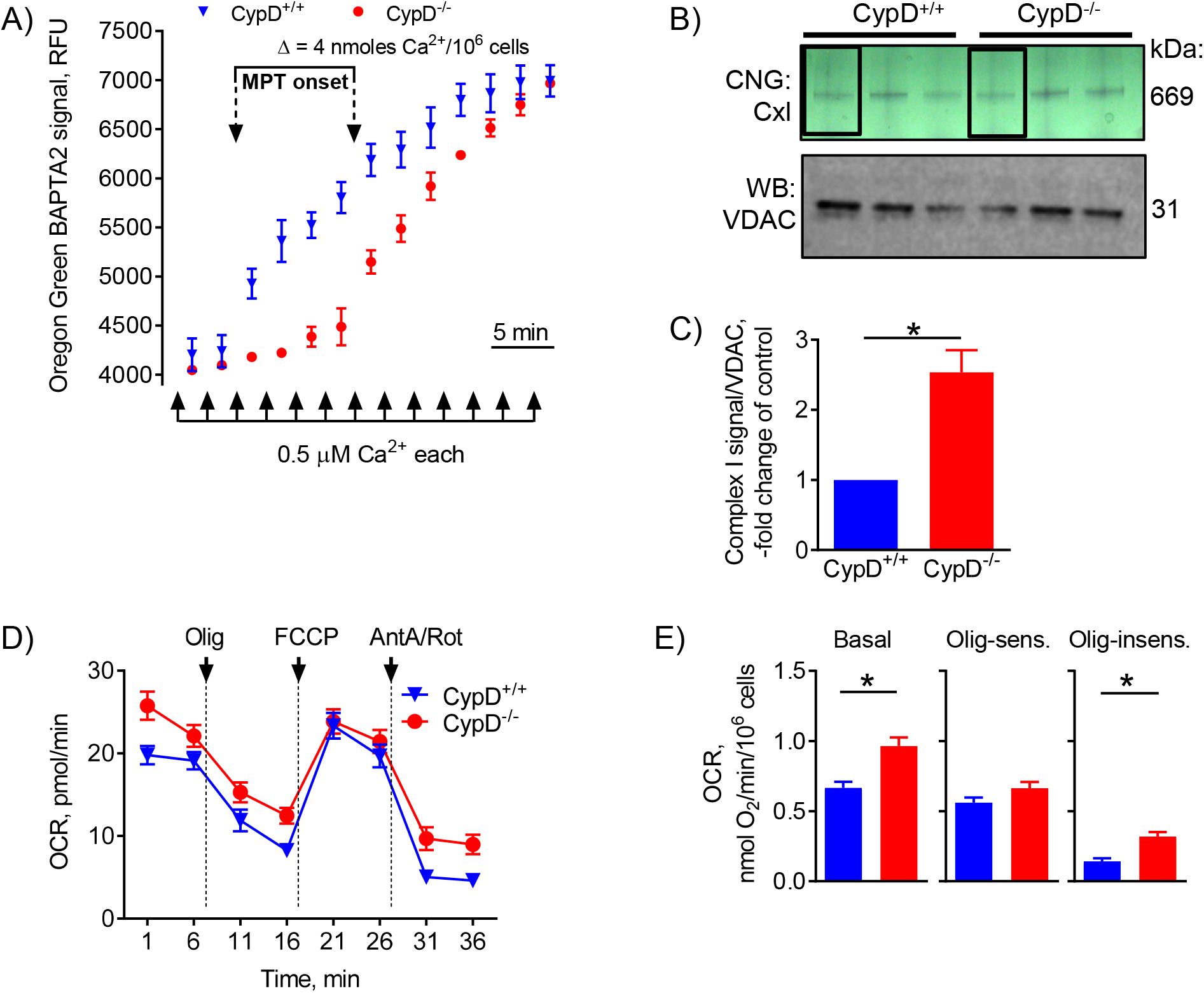
CypD KO BMSCs have more active mitochondria. A) BMSCs from CypD KO (CypD^-/-^) and WT control (CypD^+/+^) littermates were subjected to calcium retention capacity (CRC) assay which measures sensitivity to MPTP opening after excessive calcium uptake by mitochondria. CypD^-/-^ BMSC mitochondria accumulate more calcium before opening the pore and, thus show higher CRC; B) BMSCs from CypD^-/-^ and CypD^+/+^ littermates were subjected to an In-Gel Activity assay for the mitochondrial respiratory complex I (CxI). Gel is a representative of 6 Clear Native Gels (CNG). Lanes within squares are samples from undifferentiated BMSC; C) CypD^-/-^ BMSCs show a significant increase in complex I activity when compared to control CypD^+/+^ BMSCs. BMSCs were subjected to bioenergetic profiling using Seahorse XF technology. D) Oxygen consumption rate (OCR) traces before and after the additions of oligomycin (Olig), FCCP, and antimycin A/rotenone (AntA/Rot); E) Various OxPhos parameters calculated from the OCR traces. CypD^-/-^ BMSCs show a significant increase in basal and oligomycin-sensitive (ATP-linked) OCR when compared to control CypD^+/+^ BMSCs. Data are means ± SEM (n=3-6). *, *p* < 0.05, determined by unpaired *t*-test. See also Supplementary Figures S6.

In order to test mitochondrial activity directly in live cells, we performed metabolic profiling by Seahorse XF technology. Seahorse XF profiler detects cell oxygen consumption rate (OCR) as a measure of mitochondrial OxPhos activity (Fig. 6D) and extracellular acidification rate (ECAR) due to lactate production as a measure of glycolysis. CypD KO BMSCs showed significant increase in basal OCR when compared to the WT cells, consistent with CxI activity data (Fig. 6E, left panel). OCR further increased during differentiation at Day 14 but the difference between WT and CypD KO cells was no longer significant (Supplementary Fig. S6C), consistent with the hypothesis that mitochondria in undifferentiated stage CypD KO cells operate at near maximal capacity. We observed no changes in ECAR at the undifferentiated state (Supplementary Fig. S6D). It should be noted that the observed increase in OCR during osteogenic differentiation has been well documented in our lab in both primary and, thus heterogeneous BMSC cultures (4) and in more homogenous ‘BMSC-like’ cell lines, such as C3H10T1/2 (28), as well as ST2, MC3T3-E1, and DM5 cells (Eliseev lab unpublished data). Taken together these data are consistent with the established mitochondrial phenotype of the CypD KO mouse and demonstrate an increased mitochondrial activity in undifferentiated CypD KO BMSCs.

### Genetic deletion of CypD results in acetylation-sensitive increase in β-catenin activity in BMSCs

As genetic deletion of CypD promotes mitochondrial function of BMSCs, we asked whether CypD KO-mediated upregulation of mitochondrial OxPhos signals to the osteogenic program. Our recent work (28) demonstrated that activated OxPhos stimulates OB differentiation via β-catenin because mitochondria are the source of Acetyl-Coenzyme A (Ac-CoA), a substrate for β-catenin acetylation which promotes its activity, but not necessarily stability (43–46). We therefore investigated whether the β-catenin signaling pathway was affected in CypD KO BMSCs. It should first be noted that the CypD KO-mediated increase in basal OCR at Day 0 did not translate to a similarly increased ATP production as Oligomycin-sensitive ATP-linked OCR was not significantly altered between WT and CypD KO (0.56 ± 0.04 vs 0.66 ± 0.06 (nmol O_2_/min/10^6^ cells); *p* = 0.08, Fig. 6E, middle panel). As ATP production is tightly coupled to ATP demands, this may indicate that ATP demands in WT and CypD KO BMSCs at the undifferentiated stage are similarly low and the increased mitochondrial oxidative function in CypD KO BMSCs is directed towards activities other than ATP production, such as providing carbon skeletons for other cellular processes including acetylation. There was an increase in Oligomycin-insensitive OCR, also referred to as the Proton Leak, in CypD KO BMSC (Fig. 6E, right panel). This may be due to the known effect of CypD on oligomycin sensitivity of F_O_F_1_-ATPase (47), alternatively it could represent activation of alternative transport pathways which also consume the proton gradient. Conversion of mitochondrial Ac-CoA to cytosolic Ac-CoA needed for β-catenin acetylation is mediated by ATP citrate lyase (ACLY); and we previously reported that OxPhos-mediated activation of β-catenin is sensitive to inhibition of ACLY by SB204990 (SB, 28). We found that β-catenin levels in CypD KO cells were increased when compared to WT controls as measured by western blot; and this increase could be blocked using SB to inhibit ACLY mediated acetylation (Fig. 7A). These data indicate that when compared to the WT cells, CypD KO BMSCs have a small but statistically significant increase in β-catenin protein and that this increase is regulated by a mechanism dependent on Ac-CoA originated in mitochondria. As mentioned above, β-catenin acetylation supports its transcriptional activity but not necessarily stability or protein accumulation. To determine whether β-catenin activity was enhanced in an acetylation-sensitive manner in CypD KO BMSCs, we used the lentiviral TOP-GFP.mC and FOP-GFP.mC reporters that provide a β-catenin-specific and -unspecific GFP signal, respectively (48). CypD KO and WT BMSCs expressing the TOP-GFP.mC or FOP-GFP.mC lentiviral vectors were treated with SB or vehicle control to determine whether β-catenin activity was different between the studied groups and whether it was acetylation-sensitive. Cells were analyzed by flow cytometry for both ubiquitous mCherry and reporter-dependent GFP expression. As predicted, when compared to the WT controls, CypD KO BMSCs show a significant increase in β-catenin activity (GFP^TOP^ – GFP^FOP^ signal in mCherry^+^ cells) that was reversed by treatment with SB (Fig. 7B). These data indicate that CypD KO BMSCs display a bona fide increase in β-catenin activity that is sensitive to presence of Ac-CoA originated in mitochondria. RNAseq data from WT and CypD KO BMSCs further confirmed that the β-catenin pathway was indeed affected in CypD KO BMSCs, revealing a β-catenin-specific subset (49) of differentially expressed (>50% up- or down-regulated; *p* < 0.05) genes (Fig. 7C). Importantly, β-catenin target genes related to osteogenesis, e.g. *Dlx1*, *Bglap*, and *Wnt9a*, were significantly upregulated (Fig. 7C). We conclude that elevated β-catenin activity in CypD KO BMSCs, sensitive to ACLY inhibition, suggests its dependence on mitochondrial TCA cycle-derived Ac-CoA (Fig. 7D).

**Figure 7.**
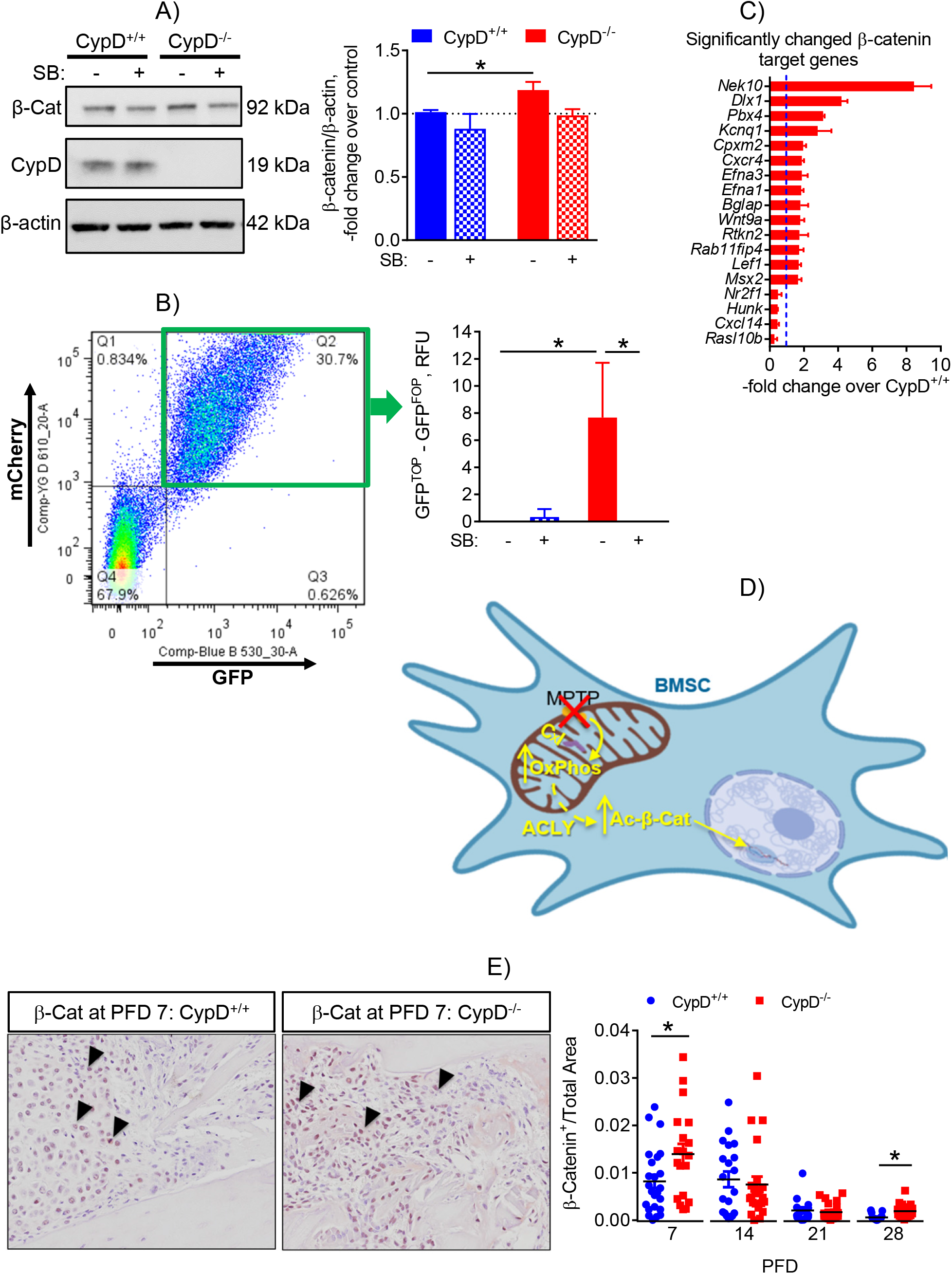
Genetic deletion of CypD results in acetylation-sensitive increases in β-catenin protein content and activity in BMSCs. CypD^-/-^ and CypD^+/+^ BMSCs were treated with either ACLY inhibitor, SB204990 (SB), or vehicle and collected for western blot (A) or flow cytometry analysis (B). In (A), blots were probed for β-catenin and then re-probed for CypD and for β-actin and analyzed with densitometry (right panel). In the graph, dotted line marks β-catenin protein level in control CypD^+/+^ cells. In (B), cells were first infected with either TOP^GFP^ β-catenin-specific or FOP^GFP^ non-specific reporters. Both reporters contained mCherry as an infection efficiency marker. Double-positive (mCherry^+^, GFP^+^) cells (B, left panel) were selected for analysis of GFP signal (B, right panel). Non-specific FOP^GFP^ signal was subtracted from TOP^GFP^ signal. CypD^-/-^ BMSCs show a significant increase in β-catenin protein levels (A) as evident from blot densitometry analysis (A, right panel) and activity (B), which was sensitive to inhibition of mitochondria-dependent acetylation with SB. RNAseq analysis showed significant changes in expression of β-catenin target genes in CypD^-/-^ vs CypD^+/+^ BMSCs (C); D) A graphical summary showing how CypD/MPTP inhibition leads to the downstream increase in acetylated β-catenin; E) Sections of calluses were probed with antibodies against β-catenin at various time-points. CypD^-/-^ mice show increased β-catenin (β-Cat) immunostaining at PFD 7 and 28. Data in are means ± SEM (n=3-4 for A – C). *, *p* < 0.05, determined by ANOVA (A & B) followed by post-hoc analysis or unpaired *t*-test (C). Plot in (E) shows actual data points and calculated means (n=8, 3 replicates each). *, *p* < 0.05, determined by unpaired *t*-test of each time point. Unpaired *t*-test was used because samples at each time point were collected and analyzed independently of other time points.

To investigate whether β-catenin is affected by CypD deletion *in vivo* during fracture healing. we performed an immunostaining for β-catenin in calluses at various time-points and observed an increase in CypD KO vs controls at PFD 7 and 28 (Fig. 7E). Of note, this increase was concomitant with the increase in osteogenic markers, osteocalcin and P1NP (Fig. 4A & B). This finding, although correlative at this stage, highlights a potential role of β-catenin as an effector of signaling caused by CypD/MPTP inhibition in fracture repair and provides a rationale for future mechanistic studies.

## Discussion

Cyclophilin D is an exclusively mitochondrially localized prolyl-peptidyl cis-trans isomerase. While the exact role for the enzymatic activity for CypD in mitochondria is not yet well understood, its position as a key regulator of the mitochondrial permeability transition pore has been repeatedly demonstrated. Both pharmacological (i.e. Cyclosporin A, NIM811, Debio025, and JW47) and genetic (CypD KO) inhibition of CypD result in mitochondria with decreased sensitivity to calcium- and reactive oxygen species-induced permeability transition, as well as increased oxygen consumption rates and increased ATP/ADP ratio. All supportive of decreased activity of the MPTP in the absence of CypD. The impact of CypD KO on cellular processes is wide spread including cardiac development (50), ischemia reperfusion injury (27), and now bone formation. Manipulation of CypD as a regulator of the MPTP, and thus mitochondrial integrity and bioenergetics is a useful mechanism for investigating cellular reliance on mitochondria-derived ATP and metabolites.

The role of mitochondrial dysfunction in bone pathologies has not been studied extensively. In our previous report, we uncovered evidence for the MPTP-mediated mitochondrial dysfunction in aging mouse bone as a cause of aging-associated bone loss (22). Building on our previous work and existing knowledge of MPTP involvement in the response to trauma and inflammation, we investigated the role of MPTP in bone fracture healing. Healing of bone fracture progresses in distinct phases: an inflammation phase that occurs acutely post fracture, soft cartilaginous callus phase, mineralization phase, and a remodeling phase. As acute inflammation involves processes linked to mitochondrially generated reactive oxygen species, and tissue repair is an energy intensive process, we hypothesized that the CypD/MPTP inhibition model would show improved fracture healing. To test this hypothesis, we used a mouse tibia fracture model. We observed that male but not female CypD KO mice had a significant increase in biomechanical properties of repaired bone at PFD 35. The increased torsional strength and rigidity of the repaired bone was associated with better bridging of the fractured cortical surfaces, increased mineralization of the bony bridge, and presence of more mature collagen, all indicative of higher OB bone forming activity during fracture healing in CypD KO mice. Histological, immunohistochemical, and serological analyses all indicated that male CypD KO mice show an accelerated bone fracture callus ossification and remodeling. An important question is why CypD KO did not show any effect on fracture healing in female mice. One possible explanation may come from the known fact that estrogen is a strong inhibitor of MPTP opening; although the mechanism of this protective effect is not yet well understood (51, 52). Therefore, because of presence of high estrogen, female mice may be well protected from MPTP opening even under the conditions of trauma such as fracture. Sexual dimorphism of MPTP activity has been reported before in various pathologies (53). These results suggest avenues for future research, such as testing the effect of CypD KO in ovariectomized or aged female mice, conditions associated with loss of estrogen.

We also used a pharmacological inhibitor of CypD, NIM811, and observed a positive effect on torsional rigidity of the repaired bone but not on maximum torque. These data suggest that further optimization of the inhibitor dose and regimen as well as developing bone-targeted variants of the inhibitor has a strong potential to enhance the inhibitor efficacy for fracture repair.

Our fracture data in male mice strongly suggest increased function of osteogenic cells during repair. As fracture repair is a complicated multicellular process and our model was not cell-specific, it is possible that the observed increase in osteogenic function is due to systemic changes. We, therefore proceeded to investigate whether MPTP loss-of-function via CypD genetic deletion or pharmacological inhibition exerts any cell-autonomous effect on BMSCs. To determine how MPTP loss-of-function via CypD KO or pharmacological inhibition affects osteogenic differentiation of BMSCs, we performed osteoinduction studies *in vitro* as well as ectopic bone formation assay *in vivo* (39). CypD KO BMSCs displayed an increase in osteogenic potential *in vitro*, along with greater ectopic bone formation *in vivo*. NIM811-mediated inhibition of CypD/MPTP also produced pro-osteogenic effect in BMSCs. Our lab and others have shown that as BMSCs undergo osteogenic differentiation, their use of mitochondrial OxPhos is upregulated and that active OxPhos, in turn, promotes osteogenic signaling (4,5,42). Herein, we demonstrate that CypD deletion and thus MPTP inactivation results in an increase in mitochondrial OxPhos activity. In addition, we show that similarly to human BMSCs and osteoprogenitors from other sources, mitochondrial capacity increases throughout the course of osteogenic differentiation in primary murine BMSCs which, to our knowledge has not been comprehensively demonstrated before. All our data thus far indicate that CypD deletion or pharmacological inhibition and subsequent MPTP closure result in more active mitochondria and exert cell-autonomous pro-osteogenic effect in BMSCs.

It has been consistently demonstrated that canonical Wnt/β-catenin signaling is activated during BMSC osteogenic commitment (28). We recently reported that mitochondrial activity in BMSCs promotes Wnt/β-catenin signaling through an increase in β-catenin acetylation during osteogenic differentiation (54). Acetylation of β-catenin is known to promote its transcriptional activity with or without the effect on protein stability (44). Thus, we wanted to determine the impact of inhibiting MPTP via CypD deletion on availability of mitochondrially generated Ac-CoA for β-catenin acetylation and thus activity. β-catenin protein levels and activity were significantly increased in CypD KO BMSCs, and this increase was sensitive to the ACLY inhibitor and, thus dependent on mitochondria-originated Ac-CoA. Thus, these data demonstrated that β-catenin is activated in acetylation-dependent manner concomitantly with the increase in mitochondrial activity in CypD KO BMSCs and could potentially be involved in this process. This observation helps explain the difference between the effect of CypD KO on *Bglap* and *Ibsp* expression in Figure 5C as *Bglap* is a known direct transcriptional target of TCF/β-catenin while *Ibsp* is not (38).

Despite the fact that the fracture model used here heals very efficiently in mice and it is, therefore hard to detect positive effects of treatments, we were able to show significant improvement in our CypD KO and NIM811-treatment models. Thus, a challenged fracture model, e.g. in aged mice, can potentially better define the role of CypD and MPTP in bone healing. The global CypD KO animal model used in our study, more closely mimics the expected action of pharmacologic CypD inhibitors lacking cell specificity and provides compelling evidence that CypD is an important therapeutic target for improving outcomes in fracture repair. Cell-specific deletion of CypD on the other hand, may provide valuable mechanistic data on the cell type and stage of differentiation at which inhibition of MPTP during fracture repair is most important and we are currently establishing Col1-Cre, Col2-Cre, and Prx1-Cre-driven deletion of CypD.

In conclusion, herein we report a novel impact of CypD/MPTP inhibition on bone fracture repair *in vivo* and osteogenic function of BMSCs *in vitro* presumably via increased mitochondrial activity supporting β-catenin signaling. We demonstrate that CypD deletion or NIM811 treatment in male mice leads to accelerated fracture callus ossification and remodeling and increased osteoblast function resulting in stronger repaired bone. Additionally, we show that stimulation of mitochondrial activity via CypD deletion or pharmacological inhibition, promotes osteogenic differentiation concomitant with an acetylation-sensitive increase in β-catenin activity *in vitro*. This work, along with previous work in our lab, provides support for the role of mitochondria, as key regulators of the osteogenic program and bone formation and repair. We posit that CypD inhibition using pharmacological inhibitors of CypD, such as NIM811, Debio025, and JW47 (16, 17), as potential candidates to induce bone formation and promote bone repair especially in aging and other pathologies.

## Acknowledgements

We would like to thank the Center for Musculoskeletal Research and the Mitochondrial Interest Group for their fruitful discussions; Frances Fuks, Alexander Chirokikh, and Julian Joseph for contributing to histomorphometry analysis; Katherine Escalera-Rivera, Alex Hollenberg, and Sarah Murphy for assistance with mouse management and prep assistance; Drs. George Porter, Jennifer Jonason and Cheryl Ackert-Bicknell for their contributions to experimental design and data interpretation; Dr. Paul Brookes for the use of Seahorse apparatus; the histology core within the Center for Musculoskeletal Research (Kathy Maltby, Sarah Mack, and Jeffery Fox), the micro-CT core within the Center for Musculoskeletal Research (Michael Thullen), and the University of Rochester Genomics and Flow Cytometry Cores. Financial support was provided by NIH (R01 AR072601 and R21 AR07928 to R.A.E.; R01 AR070613 to H.A.; and P30 AR069655 to the Center for Musculoskeletal Research) and by a grant to R.A.E. from the Orthopaedic Research and Education Foundation with additional funding provided by Musculoskeletal Transplant Foundation.

## Author Contributions

Conceptualization, R.A.E.; Investigation, B.H.S., T-J.S., K.S., R.S., A.P., G.B., L.C.S., E.G; Writing- Original Draft, B.H.S., K.S., C.O.S., E.G., E.B.; Writing- Review & Editing, B.H.S., H.A., R.A.E; Supervision, R.A.E.

## Competing Interests

The authors declare no competing interests

## Materials & Correspondence

To be addressed to Roman A. Eliseev

## Supplementary Data

**Table S1:**
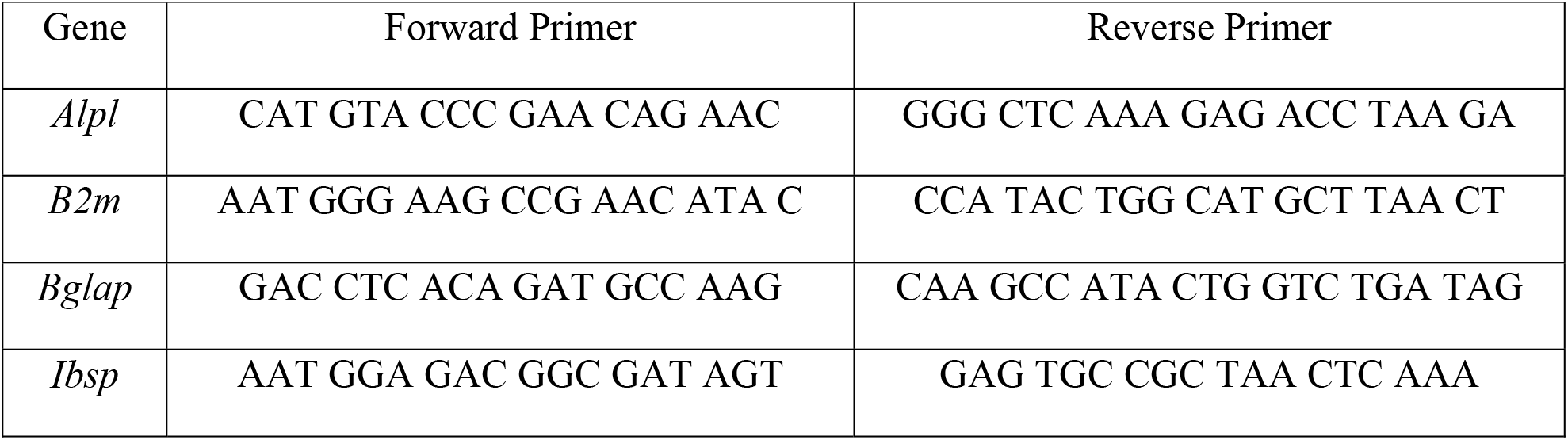
Primers used for real time RT-PCR analysis. All primers written from 5’ to 3’ (left to right).

**Table S2:** Example of power calculation used to determine the n for biomechanical testing.

| Sample Size (n) | $\sigma$ (Std Dev) | $\Delta$ (effect size) | $\alpha$ (1-p, 5%) | $\beta$ (power, 80%) |
| --- | --- | --- | --- | --- |
| 6 | 0.357 | 0.566 | 1.96 | 0.8416 |

**Supplementary Figure S1 (Related to Figure 1).**
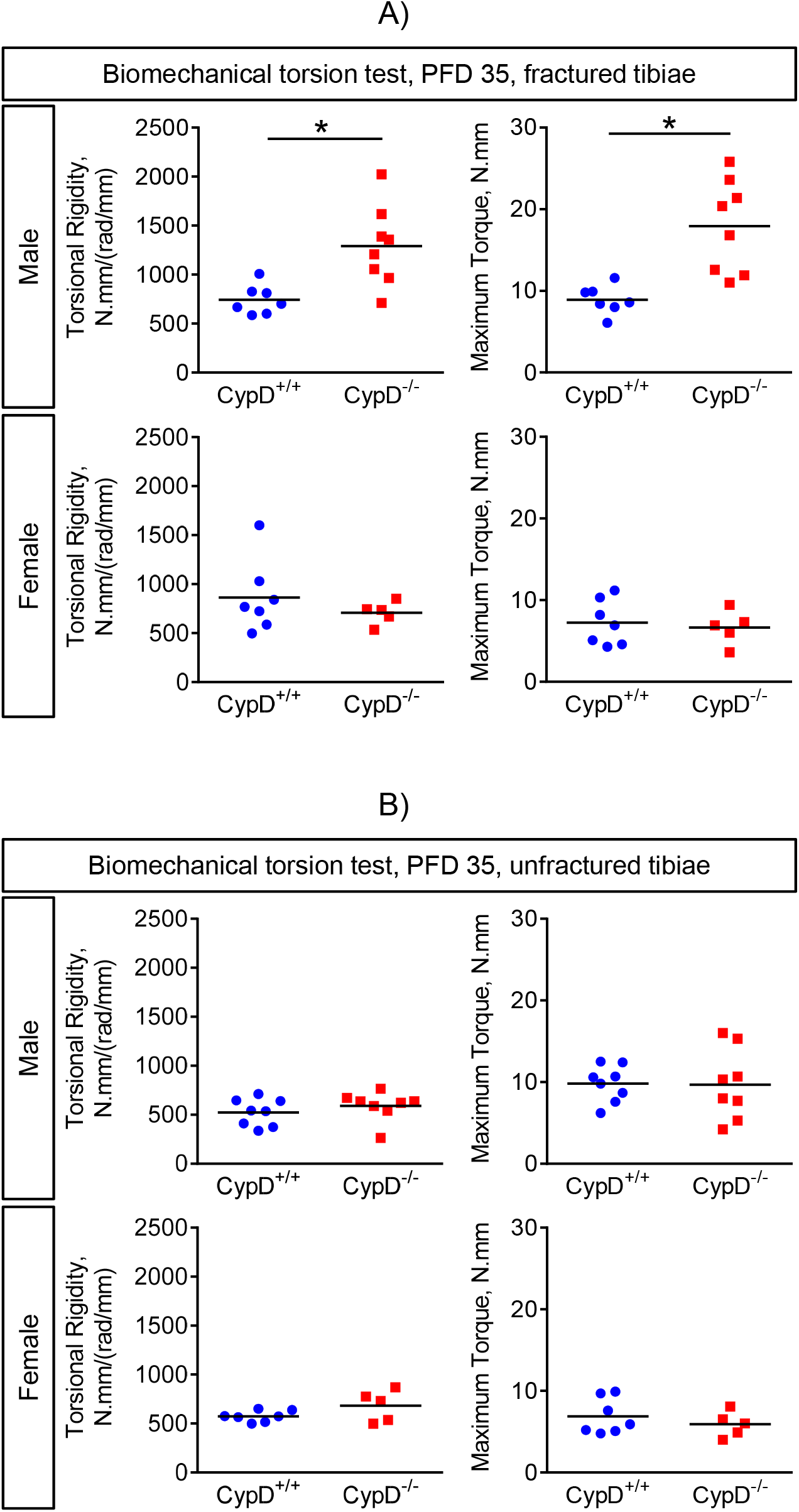
Genetic deletion of CypD results in stronger repaired bone in male mice at post fracture day 35 and no difference in unfractured bone. Tibial fractures were performed on 3-month-old male and female mice; and contralateral unfractured bones were collected for torsion testing at PFD 35. Biomechanical properties were reported as torsional rigidity indicating bone toughness and maximum torque indicating bone strength. CypD^-/-^ male but not female mice show tougher and stronger bone repair at PFD 35. No difference in torsional properties of contralateral unfractured bone when compared to control CypD^+/+^ littermates is observed in both males and females. Plots show actual data points and calculated means (n= 5-8). *, *p* < 0.05 vs CypD^+/+^ controls as determined by an unpaired *t*-test.

**Supplementary Figure S2 (Related to Figure 1).**
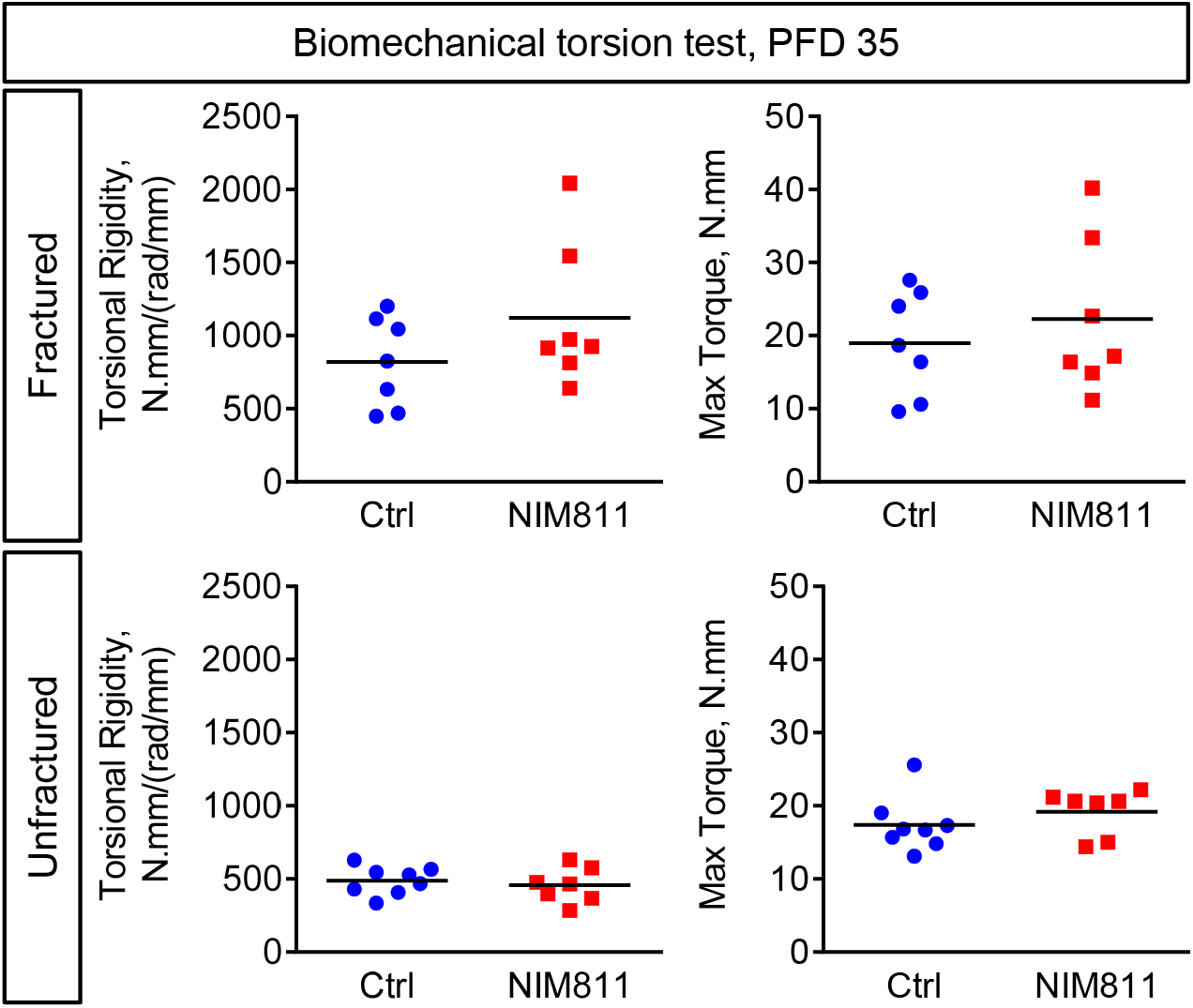
Pharmacological inhibition of CypD results in more rigid bone repair at post fracture day 35. Tibial fractures were performed on 3-month-old male and female mice; and contralateral unfractured bones were collected for torsion testing at PFD 35. Biomechanical properties were reported as torsional rigidity indicating bone toughness and maximum torque indicating bone strength. Plots show actual data points and calculated means (n=7 or 8).

**Supplementary Figure S3 (Related to Figure 1).**
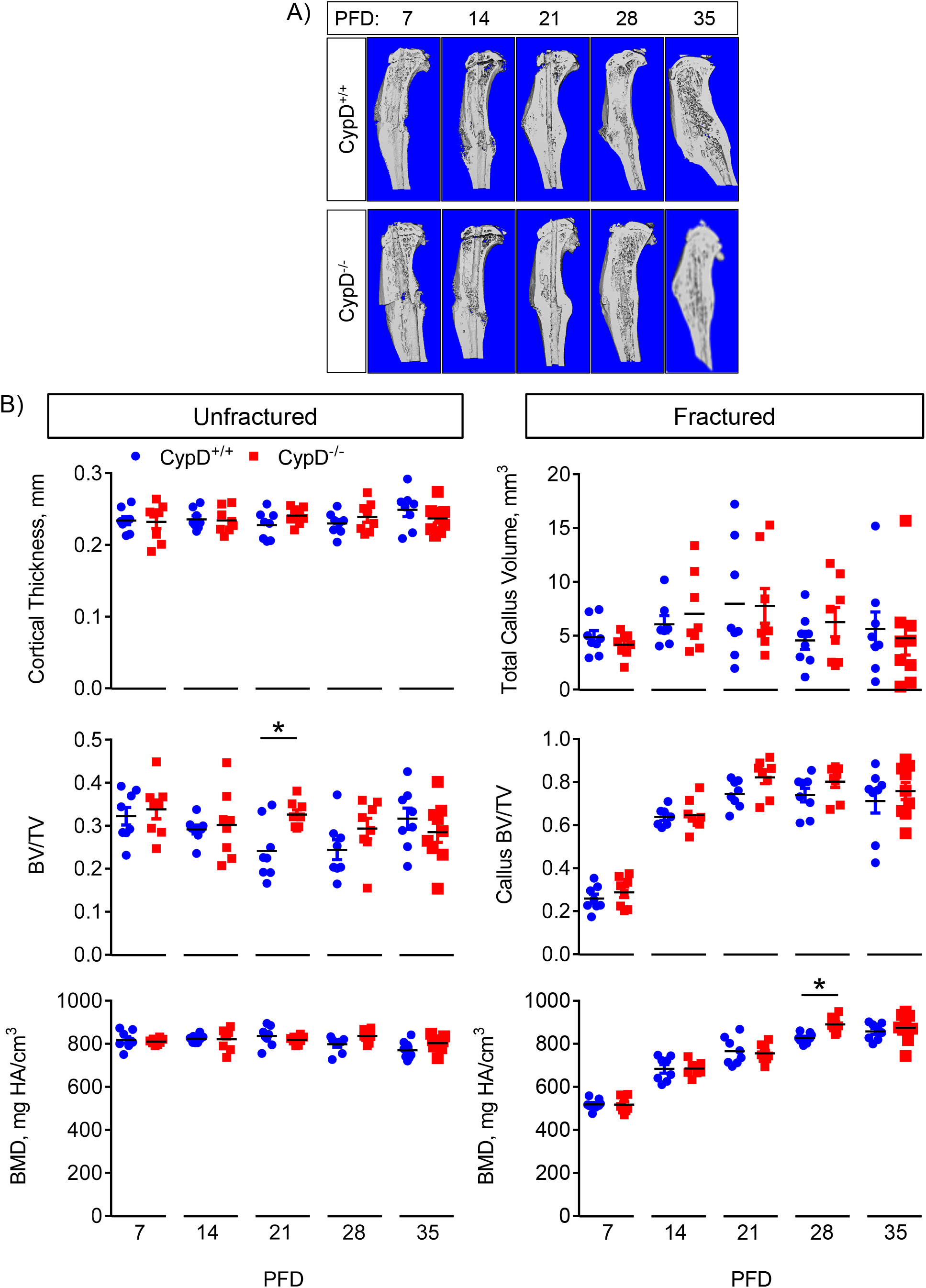
Micro-CT analysis of calluses during fracture healing. Fractured tibiae were collected at various time points and analyzed with micro-CT. Sagittal sections of the fractured tibiae are shown in A. In contralateral unfractured tibia, most of the parameters were similar in WT and CypD KO mice except for trabecular BV/TV at PFD 21 (B, middle left panel). In the fractured tibiae, most of the parameters were similar in WT and CypD KO mice apart from BMD at PFD 28 (B, bottom right panel). Plots show actual data points and calculated means (n=8). *, *p* < 0.05, determined by unpaired *t*-test of each time point. Unpaired *t*-test was used because samples at each time point were collected and analyzed independently of other time points.

**Supplementary Figure S4 (Related to Figure 4).**
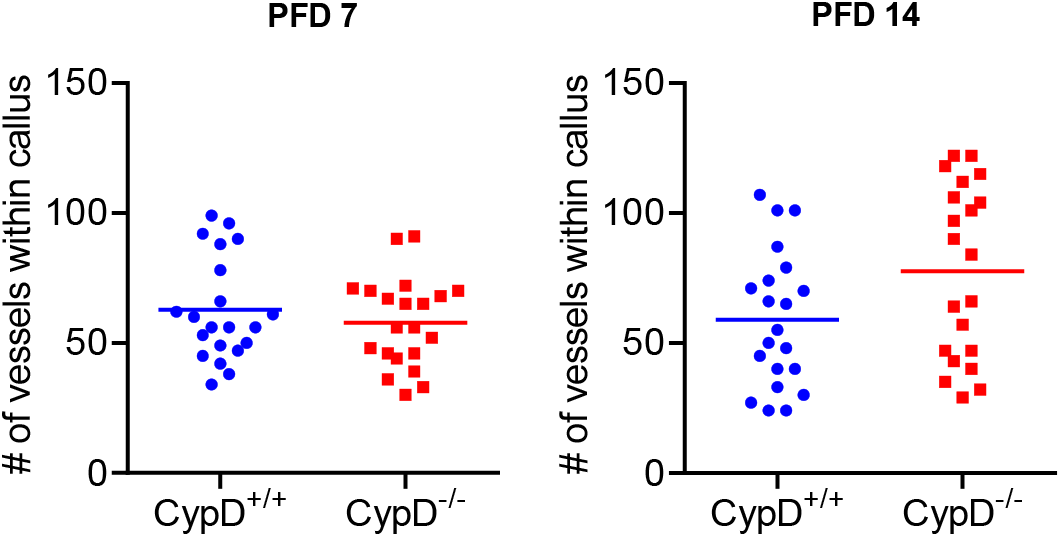
Genetic deletion of CypD does not affect vascularization at PFD 7 or 14. Fractured tibiae were collected at PFD7 and 14 and then analyzed via ABH/OG histology followed by histomorphometry analysis. CypD^-/-^ mice show no difference in vessel number at PFD7 (A) or PFD14 (B) when compared to CypD^+/+^ littermates. Plots show actual data points and calculated means (n=8).

**Supplementary Figure S5 (Related to Figures 4 & 5).**
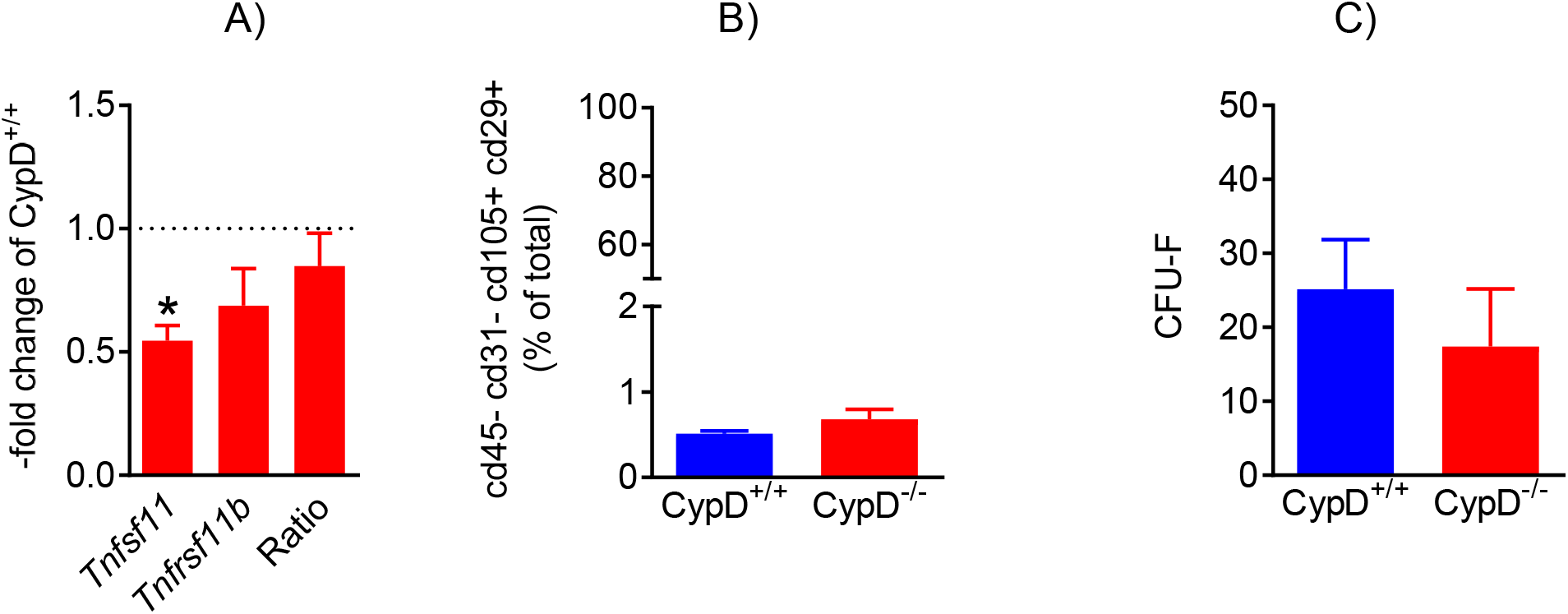
Genetic deletion of CypD does not affect BMSC population frequency, colony formation, and signaling to osteoclasts. BMSCs from CypD KO (CypD^-/-^) and WT control (CypD^+/+^) littermates were analyzed by RNAseq (A), by flow cytometry for BMSC markers (cd45- cd31- cd105+ cd29+, B), or via CFU assay (C). CypD KO BMSCS showed no difference in the ratio of RANKL gene (*Tnfsf11*) to Opg gene (*Tnfrsf11b*) expression, surface markers, or CFU-F count when compared to control BMSCs. Data are means ± SEM (n= 3-5). Statistical significance was analyzed with unpaired *t*-test.

**Supplementary Figure S6 (Related to Figure 6).**
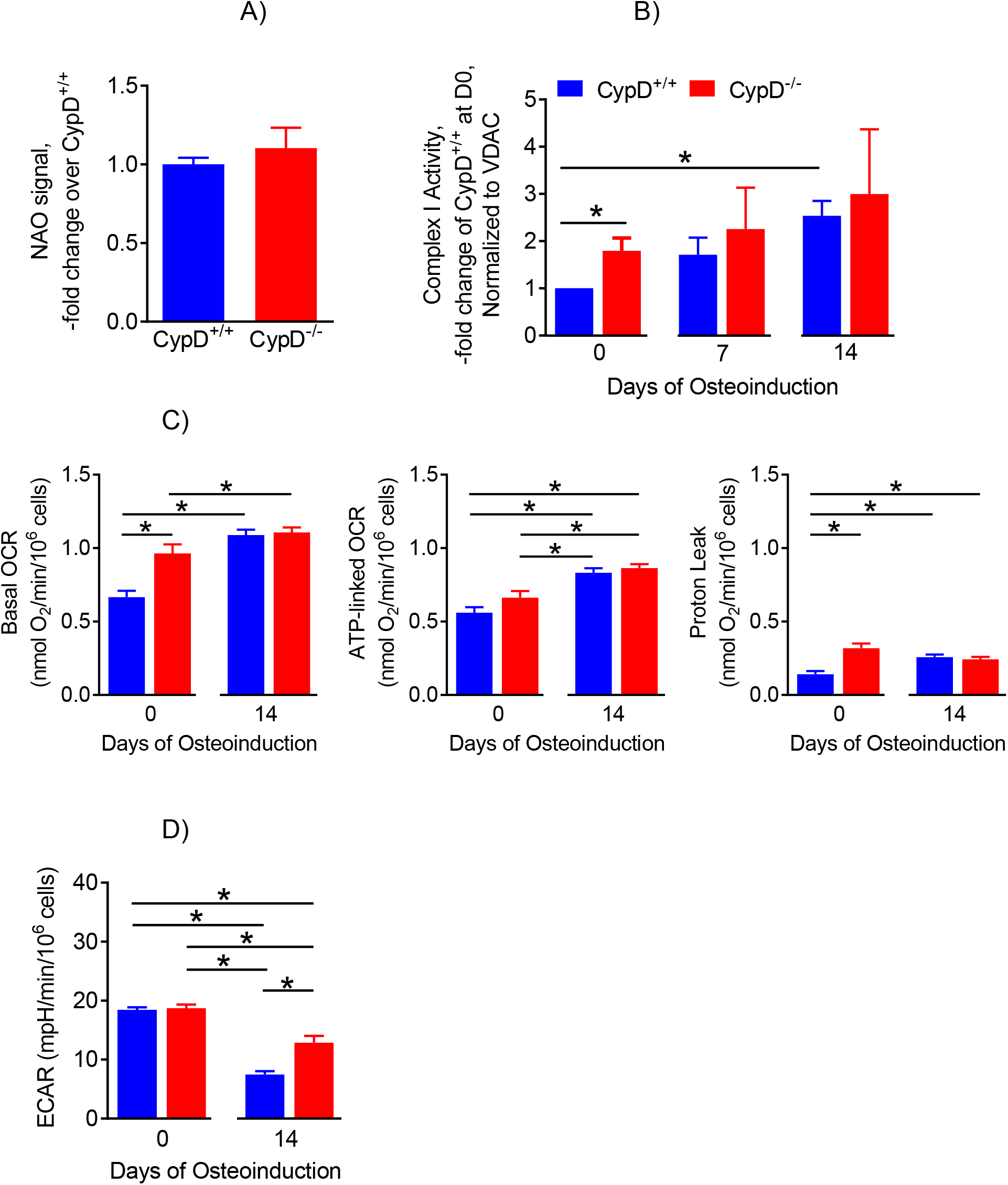
BMSC mitochondrial parameters during osteogenic differentiation. BMSCs from CypD^-/-^ and control CypD^+/+^ littermates were osteoinduced for 0, 7, and 14 days and collected for flow cytometry or subjected to in-gel activity assay for respiratory complex I or bioenergetic profiling using Seahorse XF technology. A) CypD^-/-^ and CypD^+/+^ BMSCs have similar mitochondrial mass as measured with NAO staining and flow cytometry; B) Complex I activity is similar in CypD^-/-^ and CypD^+/+^ BMSCs throughout osteogenic differentiation; C) Various mitochondrial bioenergetic parameters calculated from the inhibitory analysis; D) Extracellular Acidification Rate (ECAR) due to lactate production, a measure of glycolytic activity. Undifferentiated (D0) CypD^-/-^ BMSCs show a significant increase in basal OCR (B) and proton leak when compared to control CypD^+/+^ BMSCs. All mitochondrial parameters increased while glycolysis decreased (D) at D14 of osteoinduction in both groups of cells. Data are means ± SEM (n=4). *, *p* < 0.05, determined by ANOVA followed by post-hoc analysis.

## Notes

The authors have declared that no conflict of interest exists.

